# K29-linked unanchored polyubiquitin chains disrupt ribosome biogenesis and direct ribosomal proteins to the Intranuclear Quality control compartment (INQ)

**DOI:** 10.1101/2023.05.03.539259

**Authors:** Harsha Garadi Suresh, Eric Bonneil, Benjamin Albert, Carine Dominique, Michael Costanzo, Carles Pons, Myra Paz David Masinas, Ermira Shuteriqi, David Shore, Anthony K. Henras, Pierre Thibault, Charles Boone, Brenda J Andrews

## Abstract

Ribosome assembly requires precise coordination between the production and assembly of ribosomal components. Mutations in ribosomal proteins that inhibit the assembly process or ribosome function are often associated with Ribosomopathies, some of which are linked to defects in proteostasis. In this study, we examine the interplay between several yeast proteostasis enzymes, including deubiquitylases (DUBs), Ubp2 and Ubp14, and E3 ligases, Ufd4 and Hul5, and we explore their roles in the regulation of the cellular levels of K29-linked unanchored polyubiquitin (polyUb) chains. Accumulating K29-linked unanchored polyUb chains associate with maturing ribosomes to disrupt their assembly, activate the Ribosome assembly stress response (RASTR), and lead to the sequestration of ribosomal proteins at the Intranuclear Quality control compartment (INQ). These findings reveal the physiological relevance of INQ and provide insights into mechanisms of cellular toxicity associated with Ribosomopathies.

## Introduction

Ubiquitylation is a reversible post-translational modification involving the addition of ubiquitin (Ub) onto substrates via the activity of a cascade of Ub-activating (E1s), Ub-conjugating (E2s), and Ub ligase enzymes (E3s) [1, 2]. These enzymes first catalyze the formation of an isopeptide bond between the C-terminus of a single ubiquitin protein and a lysine residue of a substrate protein [1]. Multiple rounds of further ubiquitylation result in the formation of polyubiquitin (polyUb) chains, which are connected through isopeptide bonds between an internal lysine (K6, K11, K27, K29, K33, K48 or K63) of one Ub protein with the C-terminal glycine of another Ub protein [2]. Deubiquitylases (DUBs) are enzymes that reverse Ub chain assembly by editing or completely removing polyUb chains from substrates [3]. Free or unanchored polyUb chains that are released from substrates via DUB activity are further recycled to replenish cellular pools of monoubiquitin, which given the short half-life of Ub (∼2 hrs) [4], is important to maintain Ub homeostasis, which is particularly important when cells are subject to proteotoxic stresses such as heat shock [4–7]. Ubiquitylation regulates a variety of key biological processes, including ribosome homeostasis [8–15]. Ribosomes are highly abundant macromolecular machines that catalyze protein synthesis in all organisms and provide a platform for the initial folding, processing, and targeting of nascent polypeptide chains [16, 17]. In the budding yeast, *Saccharomyces cerevisiae*, ∼150 rRNAs and 137 ribosomal protein genes are coordinately regulated to ensure efficient ribosome assembly [18]. Unassembled or excess orphan ribosomal proteins arising from aberrant complex assembly are prone to aggregation, and they are targeted for proteasome-dependent degradation by the E3 Ub ligase, Tom1 [19–24]. Failure to degrade unassembled ribosomal proteins in yeast triggers the “Ribosome assembly stress response” (RASTR) [24, 25], which downregulates ribosomal protein gene expression by sequestering Ifh1, a master transcription factor dedicated to ribosomal protein gene expression [12], and by activating the Heat Shock Factor 1 (*HSF1*) regulon, which induces expression of several chaperones [24, 25]. In particular, *HSF1* induces the expression of *BTN2*, which encodes a small heat shock-like protein chaperone that controls the formation of an inclusion within the nucleus termed the “Intranuclear quality control compartment” (INQ). In response to genotoxic or proteotoxic stress, INQ acts as a protein quality control site for degradation of misfolded proteins and other cellular factors [26–28].

Proteomic analysis in yeast revealed that many ribosomal proteins and associated factors are ubiquitylated [29]. The relevance of this ubiquitylation for ribosome function and quality control is clear from a variety of experiments. For example, non-functional 80S ribosomes with either 18S or 25S rRNA mutations are dissociated upon ubiquitylation of ribosomal proteins by either the Mag2-Hel2-Rsp5 or Rtt101-Mms1-Crt10 E3 ligase complexes and subsequently targeted for proteasome-mediated degradation. Stalled and colliding ribosomes on translating mRNAs are also dissociated following ubiquitylation of the ribosomal proteins, Rps10 and Rps7 [29, 30]. Ubiquitylation of Rps7 is also needed for efficient general translation and for selective mRNA translation during ER stress [31] and deubiquitylation of Rps7 by Otu2 is key for release of mRNA from 40S complexes and 40S ribosome recycling to form functional 80S complexes [32]. Finally, cellular stress induced by H2O2 inhibits the deubiquitylase Ubp2, favoring the ubiquitylated state of elongating ribosomes, and inhibiting translation elongation [15, 33].

While roles for both monoUb and polyUb modification in the degradation of ribosomal proteins or translation control are clear [23, 34, 35], potential roles for unanchored (substrate free) polyUb chains on ribosome biology remain unexplored [1, 36–38]. Conventional K48- and K63-linked unanchored polyUb chains are involved in protein kinase regulation [36, 39] and activate the deacetylase HDAC6 to promote the clearance of protein aggregates [40], suggesting that other unanchored polyUb chains may have significant cellular roles. However, the functions of unconventional unanchored polyUb chains, such as K6, K11, K27, K33 and K29 chains, remain to be discovered [36, 41].

One strategy for studying the physiological consequences of unanchored polyUb chains would involve assessing their abundance and influence in cells lacking DUB function [3, 36]. Because there are ∼100 different human DUB genes [42, 43], this approach is complicated in human cells, but is more tractable in yeast which expresses 22 DUB genes. In this study, we tested all possible DUB gene pairs for genetic interactions and confirmed the results of a large-scale assessment of genome-wide genetic interactions [44, 45]. One double mutant strain lacking two DUBs, *UBP2* and *UBP14*, exhibited a severe growth defect and accumulated K29-linked unanchored polyUb chains. These K29-linked unanchored chains associated with maturing ribosomes and induced RASTR, suggesting that *UBP2* and *UBP14* are important for normal ribosome biogenesis through their negative regulation of K29-linked unanchored polyUb chain levels. We used proteome-wide imaging screens to show that activation of RASTR led to INQ formation and sequestration of a subset of ribosomal proteins. Our study illustrates an intricate functional connection between ubiquitin homeostasis and ribosome assembly, defines a physiological function for INQ and provides insights into the potential mechanisms of cellular toxicity associated with mutations observed in Ribosomopathies.

## Results

### Ubp2 and Ubp14 are genetically redundant

Previous work from our laboratory mapped a complete genetic network for yeast, which involved tests of all possible gene pairs for genetic interactions, which occur when a double-mutant strain shows a growth phenotype not predicted based on the single mutant phenotypes [44–46]. The network includes tests of all pairs of 22 yeast DUB genes and revealed a negative genetic interaction involving one gene pair, *UBP2* and *UBP14* [44, 45] (Figure S1A-B). We confirmed a strong negative genetic interaction between *UBP2* and *UBP14*, where the double mutant had a severe growth defect not seen in either single mutant alone (Figure 1A). The *ubp2Δ ubp14Δ* growth defect was exacerbated at high temperature, which activates proteostasis, suggesting *UBP2* and *UBP14* share a critical cellular role, particularly during heat stress (Figure 1A). Consistent with a shared role in deubiquitylation, *ubp2Δ ubp14Δ* double mutant cells accumulated high molecular weight ubiquitylated protein species that appear to be made up of a series of roughly equidistant ubiquitin bands (Figure 1B).

**Figure 1:**
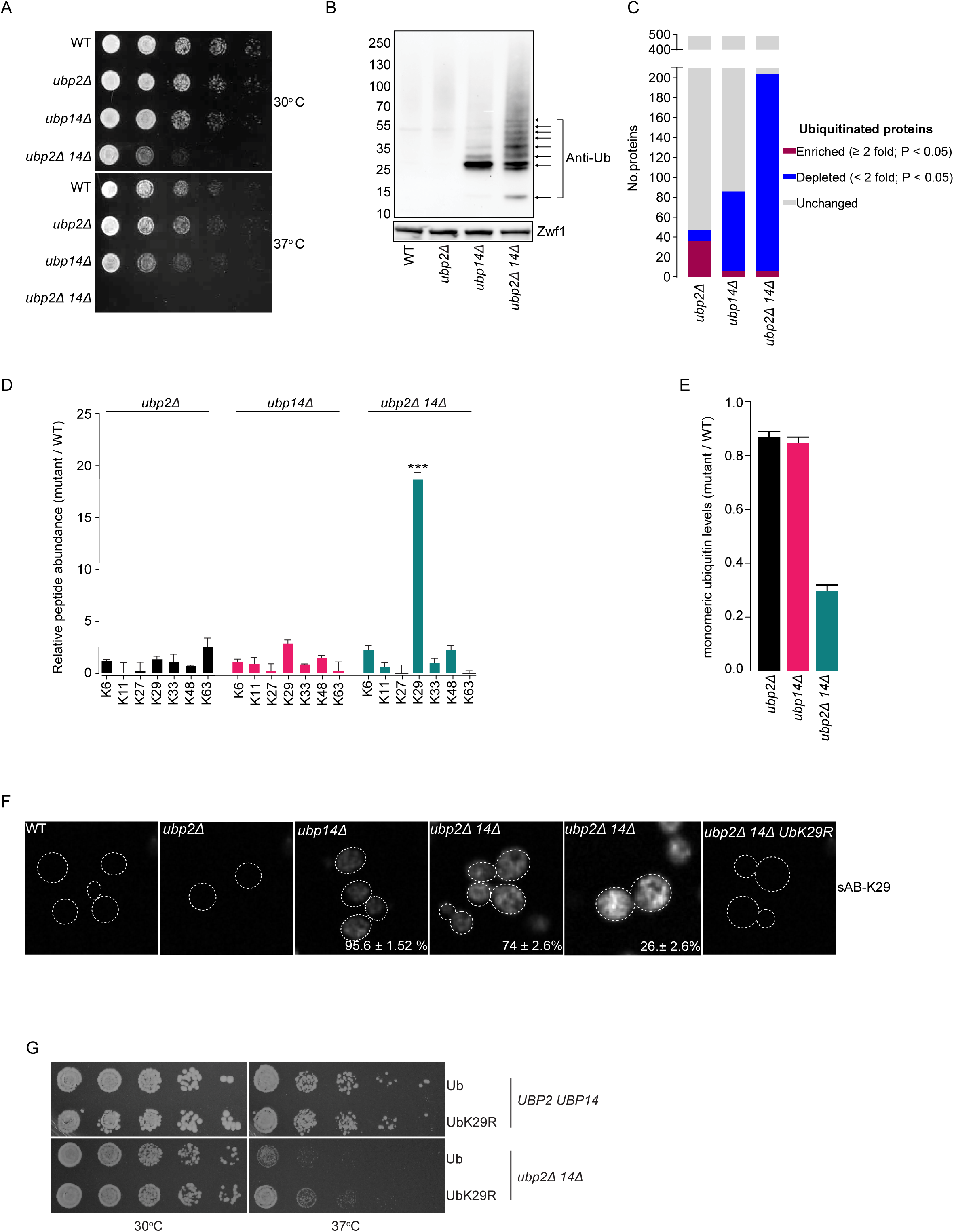
Redundant DUBs Ubp2 and Ubp14 recycle K29-linked unanchored polyUb chains. (A) Serial spot dilutions of *ubp2*Δ and *ubp14*Δ single and double mutants. The indicated strains were plated onto minimal medium and incubated at 30°C or 37°C for 1 day (WT= wild type). (B) Immunoblot of the ubiquitinome isolated from extracts of the indicated yeast strains using Tandem Ubiquitin Binding Entity (TUBE) affinity matrix-based pull downs. Arrows indicate equidistant prominent bands of ubiquitin. Molecular weight markers are shown on the left, and the lower panel shows a loading control (anti-Zwf1). (C) Label-free quantitative mass spectrometry on ubiquitylated peptides isolated using Lys-ℇ-Gly- Gly (diGly) antibodies (n=3). Fraction of proteins that are enriched (≥2-fold, P < 0.05; blue) or depleted (<2-fold, P < 0.05; red) for ubiquitin modification in *ubp2Δ*, *ubp14Δ* or *ubp2Δ 14Δ* mutant cells relative to *wild type* cells are shown. The fraction of proteins whose ubiquitination status remained unchanged in the indicated mutant relative to wild type is shown in grey. (D) Graph summarizing the relative number of 7 different polyUb chains quantified in the indicated DUB mutants using label-free quantitative Lys-ℇ-Gly-Gly (diGly) proteomics mass spectrometry (n=3). Error bars indicate standard deviation (n=3) and *** = p<0.001. (E) The relative amount of free monomeric ubiquitin in the indicated mutants measured by immunoblots with anti-Ub antibodies (n=3). Y-axis: Wild type free monomeric ubiquitin levels is set to 1.0. Error bars indicate standard deviation (n=3) and * = p < 0.05. (F) Immunofluorescence based staining of K29 polyUb chains using sAB-K29 [50] in wild type (WT) and the indicated mutant strains. For each of three biological replicates, one hundred cells of the indicated genotypes were counted to calculate the percentage of cells with the depicted phenotypes. Standard deviations are indicated (n=3). (G) Serial spot dilutions of a wild type strain (WT) or a *ubp2Δ 14Δ* mutant expressing either wild type (Ub) or a UbK29R mutant derivative as the sole source of ubiquitin. Plates were incubated at the indicated temperatures and imaged after 48 hours.

### Ubp2 and Ubp14 recycle K29-linked unanchored polyUb chains

*UBP14* is known to play a role in recycling unanchored polyUb chains [5], although the functional consequences of unanchored polyUb chain accumulation are unknown [36, 47, 48]. Moreover, a *ubp14Δ* mutant does not exhibit significant growth defects at 30 °C or 37 °C, indicating that either accumulation of free polyUb chains does not affect yeast fitness or that their levels are not high enough to compromise cell function due to existence of a redundant DUB, such as *UBP2*, which can also regulate free polyUb chain levels [5].

To explore these ideas, we utilized Lys-ℇ-Gly-Gly (diGly) antibodies to immunoprecipitate ubiquitylated peptides from cell extracts derived from wild type cells, *ubp2Δ* and *ubp14Δ* single mutant cells, and *ubp2Δ ubp14Δ* double mutant cells. Ubiquitylated proteins were quantified using label-free mass spectrometry, and we identified the fraction of proteins that were enriched or depleted for Ub modification in *ubp2Δ*, *ubp14Δ* or *ubp2Δ ubp14Δ* mutant cells relative to wild type cells (Figure 1C, Table S1). Intriguingly, while an increase in high molecular weight ubiquitylated species was observed in *ubp2Δ ubp14Δ* double mutant cells (Figure 1B), diGly proteomics identified relatively few proteins that were significantly ubiquitylated in the double mutant cells relative to wild type cells (Figure 1C). These observations suggest that *ubp2Δ ubp14Δ* double mutant cells may accumulate free polyUb chains rather than ubiquitylated substrate proteins. Indeed, the *ubp2Δ ubp14Δ* double mutant displayed a dramatic and specific increase in K29-linked polyUb chains (Figure 1D). Concomitantly, we observed a decrease in free ubiquitin pools (Figure 1E), suggesting a potential role for Ubp2 and Ubp14 in recycling K29-linked unanchored polyUb chains. The longer polyUb chains accumulate only in the absence of both Ubp2 and Ubp14, suggesting that both DUBs can recycle polyUb chains to generate smaller chains. However, we observed obvious accumulation of smaller chains in the *ubp14Δ* single mutant may indicate that Ubp2 is less effective on smaller Ub chains *in vivo* (Figure 1B).

Consistent with the recently described propensity of polyUb chains to form amyloid-like structures *in vitro* and *in vivo* [49], immunofluorescence analysis with an antibody specific for K29-linked polyUb chains (sAB-K29) [50] revealed an accumulation of fibrous network-like structures in ∼74% and bright-dense fibrillar aggregates in ∼26% of *ubp2Δ ubp14Δ* cells (Figure 1F). Similar, albeit significantly weaker, fibrous network-like structures (but no dense bright aggregates) were observed in ∼96% of *ubp14Δ* cells but not in a *ubp2Δ* or wild type cells (Figure 1F), providing further evidence to support a redundant role for *UPB14* and *UPB2* in maintaining physiological levels of K29-linked polyUb chains.

### The *ubp2Δ ubp14Δ* growth defect reflects accumulation of K29-linked unanchored polyUb chains

To more concretely link K29 linkages to the formation of the observed fibrous aggregate-like structures, we constructed a *ubp2Δ ubp14Δ* double mutant strain that carried a Ub variant in which K29 was replaced with an arginine residue (UbK29R). We failed to observe fibrous aggregate-like structures in a *ubp2Δ ubp14Δ* double mutant cells expressing UbK29R as the sole source of Ub (Figure 1F). Moreover, the UbK29R mutant partially suppressed the *ubp2Δ ubp14Δ* double mutant growth defect (Figure 1G). Thus, accumulation of K29-linked unanchored polyUb chains appears to adversely affect growth of a *ubp2Δ ubp14Δ* mutant strain.

Given that K29-linked chains are toxic to *ubp2Δ ubp14Δ* cells, we reasoned that deletion of the cognate E3 ligase gene responsible for assembly of K29-linked unanchored polyUb chains should suppress the temperature sensitive (TS) growth defect of a *ubp2Δ ubp14Δ* double mutant. To test this possibility, we constructed a conditional double mutant by engineering *ubp14Δ* cells with a *MAL32*pr-*UBP2* gene, in which *UBP2* expression was controlled by a glucose-repressible but maltose-inducible promoter [51]. The resulting *MAL32*pr-*UBP2 ubp14Δ* strain exhibited a clear fitness defect when grown on a glucose carbon source (Figure S2A). Using Synthetic Genetic Array (SGA) methodology [52], we crossed the *MAL32*pr-*UBP2 ubp14Δ* query strain to a selected array of single mutant strains, each deleted for a gene involved in the ubiquitin proteasome system (Table S2) [37, 53]. In total, we analyzed ∼100 haploid triple mutants and identified 8 deletion mutant alleles that suppressed the TS growth defect of the *MAL32*pr-*UBP2 ubp14Δ* query strain grown on a glucose carbon source. These suppressor genes included the quality control ubiquitin ligases *SAN1* (E3), *UFD2* (E4), *UFD4* (E3) and *HUL5* (E3), as well as the heat shock proteins, *APJ1*(Hsp40), *BTN2* (sHsp), *HSP42* (sHsp) and *HSP104* (AAA+ chaperone) and their suppression was confirmed by comparing the growth of *ubp2Δ ubp14Δ* cells to the triple mutant strains using a plate-based growth assay (Figure 2A).

**Figure 2:**
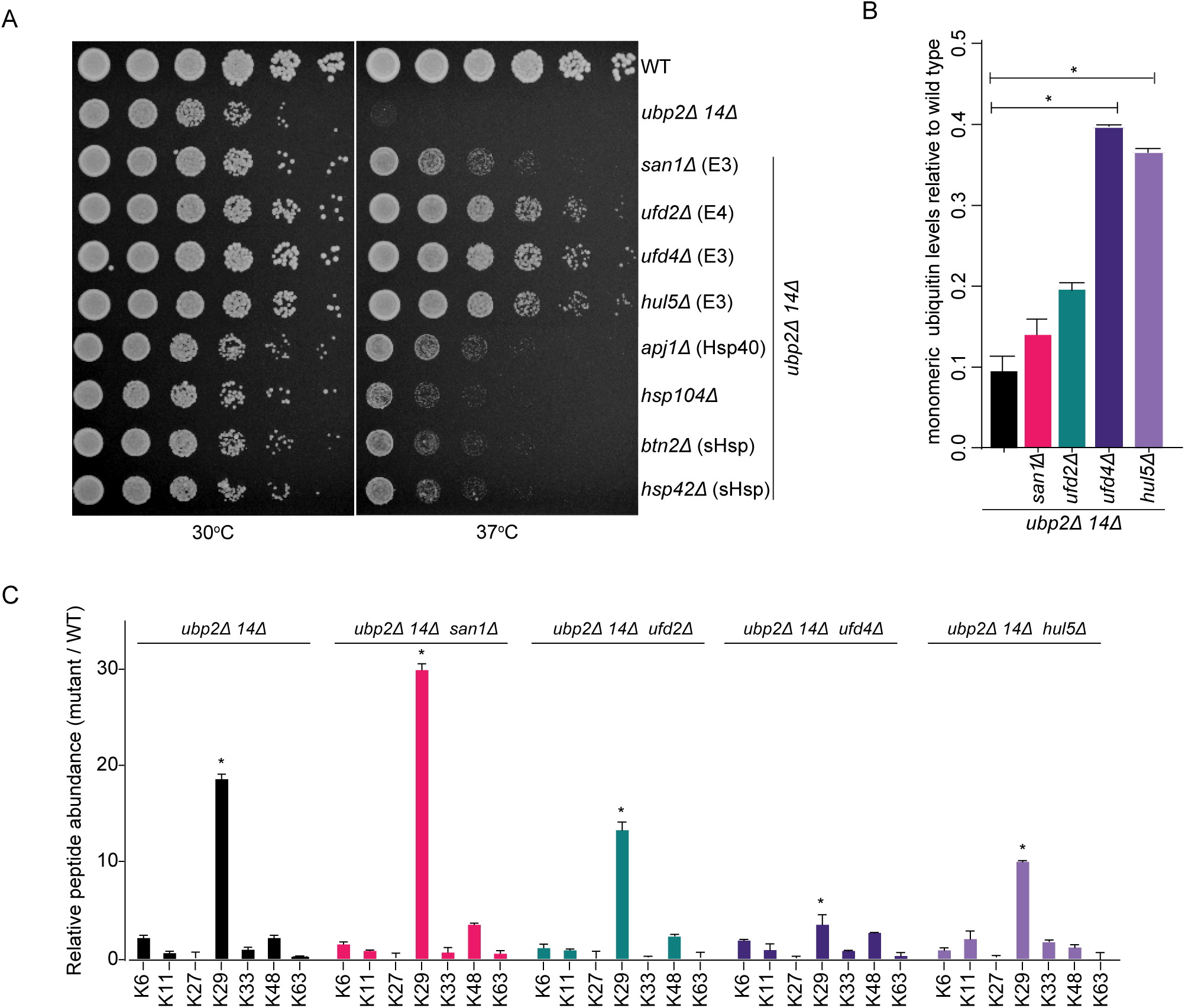
Ufd4 and Hul5 E3 ubiquitin ligases are involved in the synthesis of K29-linked unanchored polyubiquitin chains. (A) Serial spot dilutions wild-type (WT), a *ubp2Δ 14Δ* double mutant (top two rows) and *ubp2Δ 14Δ* double mutant carrying deletion of the indicated genes. Plates were incubated at 30°C or 37°C as indicated. E3= E3 ligase; E4=E4 ligase; sHsp – small Heat Shock Protein. (B) The relative amount of free monomeric ubiquitin in the indicated mutants measured by immunoblots using anti-Ub antibodies (n=3; * = P<0.05). Error bars indicate standard deviations. (C) Graph summarizing the relative amount of 7 different polyUb chains quantified in the indicated yeast mutant strains using label-free quantitative Lys-ℇ-Gly-Gly (diGly) proteomics mass spectrometry (n=3; * = P < 0.05). Error bars indicate standard deviations.

Deletion of *UFD2*, *UFD4*, or *HUL5* was associated with an increase in the number of ubiquitylated peptides (Figure S2B) and the free Ub pool (Figure 2B), with a concomitant decrease in K29- linked polyUb levels in the *ubp2Δ ubp14Δ* double mutant background (Figure 2C), with these effects being most obvious in the *ufd4*Δ and *hul5*Δ strains (Figures 2B, S2B). Thus, two E3 ligases, Ufd4 and Hul5, contribute significantly to K29-linked unanchored polyUb chain levels. We note that although the levels of free Ub increased in *ubp2Δ ubp14Δ* double mutant cells when either *UFD4* or *HUL5* were deleted, they were still significantly lower than the levels of free Ub observed in wild type cells, which suggests that the TS phenotype of a *ubp2Δ ubp14Δ* double mutant reflects accumulation of K29-linked unanchored polyUb chains rather than depletion of the free Ub pool. This is consistent with the fact that loss of *UFD4* but not Ub over-expression suppressed the growth defect of the *ubp2Δ ubp14Δ* double mutant (Figure S2C). Interestingly, unlike *UFD4* and *HUL5*, loss of *SAN1* in the *ubp2Δ ubp14Δ* double mutant dramatically increased the levels of K29-linked polyUb chains (Figure 2C), potentially implicating a role for San1 in turnover of these chains. Both *SAN1* and *APJ1*, whose loss leads to a comparable degree of suppression of the TS phenotype of a *ubp2Δ ubp14Δ* double mutant (Figure 2A), are implicated in degradation of misfolded proteins including those localized to INQ [26, 54]. Therefore, in this case, the mechanism of suppression may be indirect through stabilization of essential cellular factors aggregating in the nucleus.

### Proteome-wide imaging screens reveal *ubp2Δ ubp14Δ* double mutant defects in protein abundance and inclusion formation

To further characterize the functional impact of K29-linked unanchored polyUb chain accumulation at the level of the proteome, we made use of a single-cell imaging pipeline that exploits the yeast protein-GFP collection, a resource consisting of thousands of strains expressing different proteins tagged C-terminally with GFP, at their endogenous loci [55–57]. We used the Synthetic Genetic Array (SGA) method to cross the global GFP protein fusion library [55] into *ubp2Δ*, *ubp14Δ* and *ubp2Δ ubp14Δ* mutants, and we examined the recombinant progeny for changes in protein localization and abundance using both automated image analysis [57] and visual inspection. We focused on ∼250 proteins that reproducibly localized to punctate structures specifically in *ubp2Δ ubp14Δ* double mutant cells, but not in the corresponding single mutant or wild-type cells (Table S3). Strikingly, we identified all 26 proteins previously reported to aggregate in the Intranuclear Quality Control (INQ) compartment, which localizes within the nucleus and is positioned adjacent to the nucleolus and diagonally opposite to the spindle pole body [26, 27] (Figure 3A).

**Figure 3:**
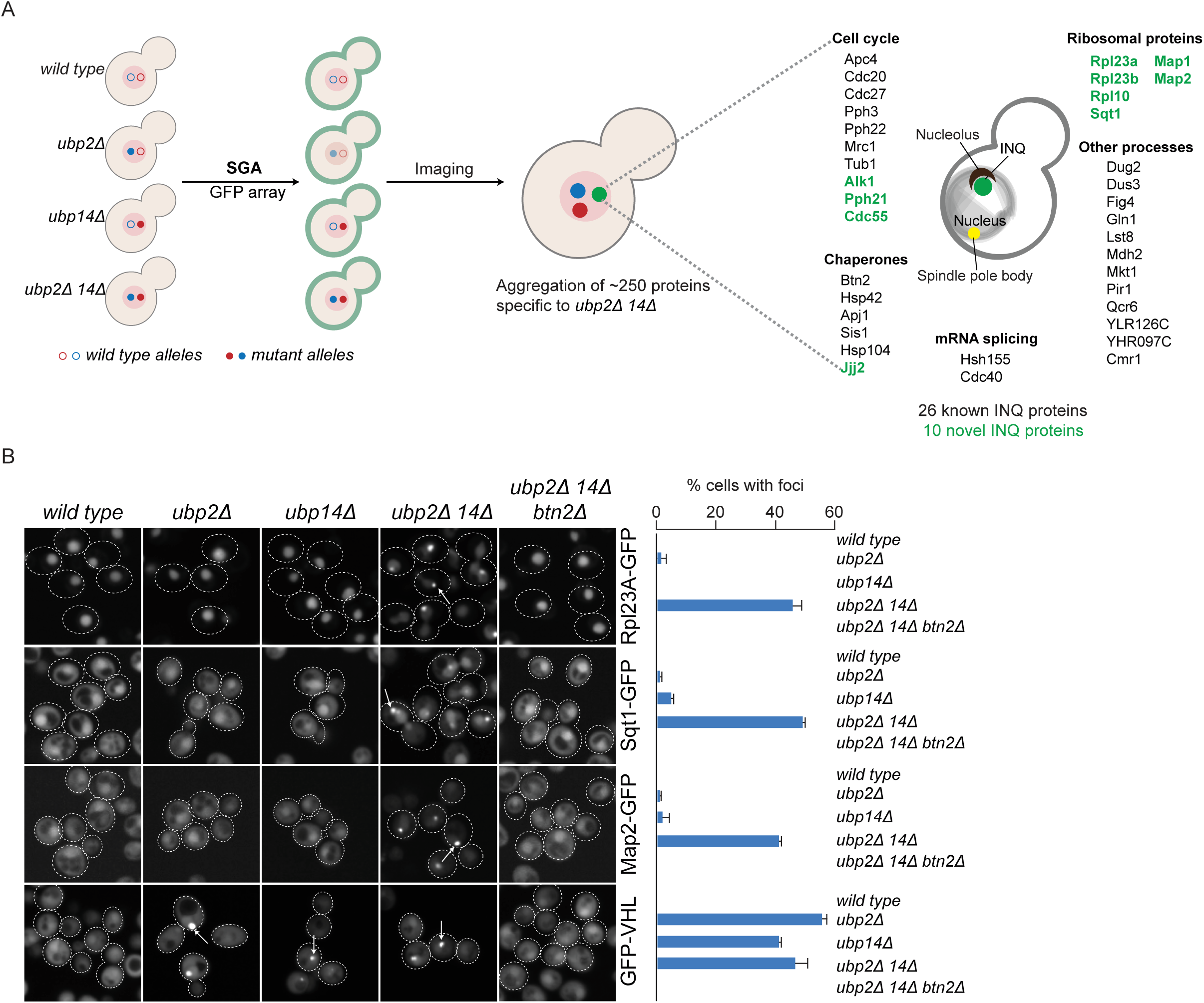
Proteome-wide imaging screens reveal *ubp2Δ ubp14Δ-*dependent defects in ribosome homeostasis. (A) Schematic depicting high-throughput imaging of the yeast GFP library [55] in *wild type*, *ubp2Δ*, *ubp14Δ* and *ubp2Δ ubp14Δ* double mutant cells. Previously identified (black) and novel (green) protein aggregates that accumulate at the Intranuclear quality control site (INQ) are highlighted. The spindle pole body (yellow), nucleolus (brown) and INQ (green) are depicted on the cell cartoon. (B) Example micrographs of yeast cells expressing specific GFP-fusion proteins in the indicated mutant strains. Arrows indicate GFP-protein aggregates. The percentage of cells with GFP-fusion protein inclusions, including Rpl23A-GFP, Sqt1-GFP, Map2-GFP, GFP-VHL, in the indicated mutant strains is shown (right panel). Error bars indicate standard deviation (n=2).

To determine which punctate structures specifically corresponded to INQ, we re-examined the localization of the ∼250 aggregated proteins in a *ubp2Δ ubp14Δ* strain that was also deleted for *BTN2*, a chaperone required for INQ formation [26, 58]. In total, 36 proteins failed to form inclusions in *ubp2Δ ubp14Δ btn2Δ* triple mutant cells, including the 26 proteins previously shown to localize to INQ, suggesting that we identified 10 novel INQ components. GFP-VHL, a constitutively misfolded model protein substrate that localizes to INQ under proteotoxic stress [26-28, 58, 59], also accumulated at INQ in a *ubp2Δ ubp14Δ* double mutant. Indeed, GFP-VHL formed aggregates in *ubp2Δ, ubp14Δ,* and *ubp2Δ ubp14Δ* mutants in a Btn2-dependent manner (Figure 3B), further indicating that loss of *UBP2* and *UBP14* can lead to protein aggregation within the INQ nuclear compartment. Interestingly, while GFP-VHL INQ aggregates segregated to mother cells during cell division in *ubp2Δ* and *ubp14Δ* single mutants (Movie S1-S2), ∼25 % of GFP-VHL aggregates segregated to daughter cells in a *ubp2Δ ubp14Δ* double mutant (Movie S3). This observation suggests that *UBP2* and *UBP14* are involved in the asymmetric segregation of INQ- localized aggregates to mother cells, a hallmark of mother cell replicative ageing and daughter cell rejuvenation [60, 61].

### Loss of *UPB2* and *UBP14* results in ribosome biogenesis defects

The ∼250 proteins found in punctate structures in the *ubp2Δ ubp14Δ* mutant were enriched for proteins involved in translation-related functions (Table S3), and 6/10 of the novel INQ components were ribosomal proteins (Rpl23A, Rpl23B, Rpl10), a ribosomal protein chaperone (Sqt1), or translation factors, Map1 and Map2, which are methionine amino peptidases that catalyze the cotranslational removal of the N-terminal methionine from nascent polypeptides (Figure 3A) [16, 18]. We note that Ub-Rps31 and Ub-Rpl40, specialized Ub-fusion ribosomal proteins, were processed normally in a *ubp2Δ ubp14Δ* double mutant, indicating that ribosomal protein aggregation was not due to defective cleavage of Ub from these proteins (Figure S3A, B) [62–64]. Moreover, cycloheximide chase experiments showed that the stability of at least one of these proteins, Sqt1, was decreased in a *ubp2Δ ubp14Δ* double mutant, suggesting that ribosomal proteins that accumulate at INQ may be subject to degradation (Figure S3C). However, the constitutive degradation of the highly unstable GFP-VHL reporter, which is independent of its localization to INQ [59, 65], was not affected in the *ubp2Δ ubp14Δ* double mutant, suggesting that depletion of the free ubiquitin pool and accumulation of K29-linked unanchored polyUb chains in *ubp2Δ ubp14Δ* cells does not affect protein degradation in general (Figure S3C).

Consistent with a role in ribosome biogenesis, polysome profiling revealed that a *ubp2Δ ubp14Δ* double mutant had decreased levels of 60S, 80S subunits and polysomes (Figure 4A) and measurements of the levels of rRNA precursors showed a decrease in intensity of the 27SA2 band relative to 35S rRNA band indicating a defect in rRNA processing (Figure S4A). Also, many ribosomal and ribosome biogenesis (RiBi) proteins were mislocalized and aggregated in the nucleolus, nucleus and cytosol in the *ubp2Δ ubp14Δ* double mutant (Figure S4B; Table S3), a strong indicator of a defect in ribosome biogenesis[18, 21, 22, 25, 66–68]. Furthermore, we also observed that the master regulator of ribosomal protein gene expression, Ifh1 [24], localized to*<ι> </i>*punctate structures in a *ubp2Δ ubp14Δ* mutant (Figure 4B), suggesting that the Ribosome Assembly Stress Response (RASTR) was active in these cells [24, 25]. RASTR is characterized by increased expression of *HSF1* regulon genes with the concomitant downregulation of genes encoding ribosomal proteins [24, 25]. Indeed, chromatin immunoprecipitation (ChIP-Seq) assays, using Rpb1 to monitor RNA polymerase II (RNAPII) gene occupancy, showed that RASTR was more strongly induced in *ubp2Δ ubp14Δ* double mutant cells compared to either of the corresponding *ubp2Δ* or *ubp14Δ* single mutants (Figure 4C, Table S4).*<ι>*

**Figure 4:**
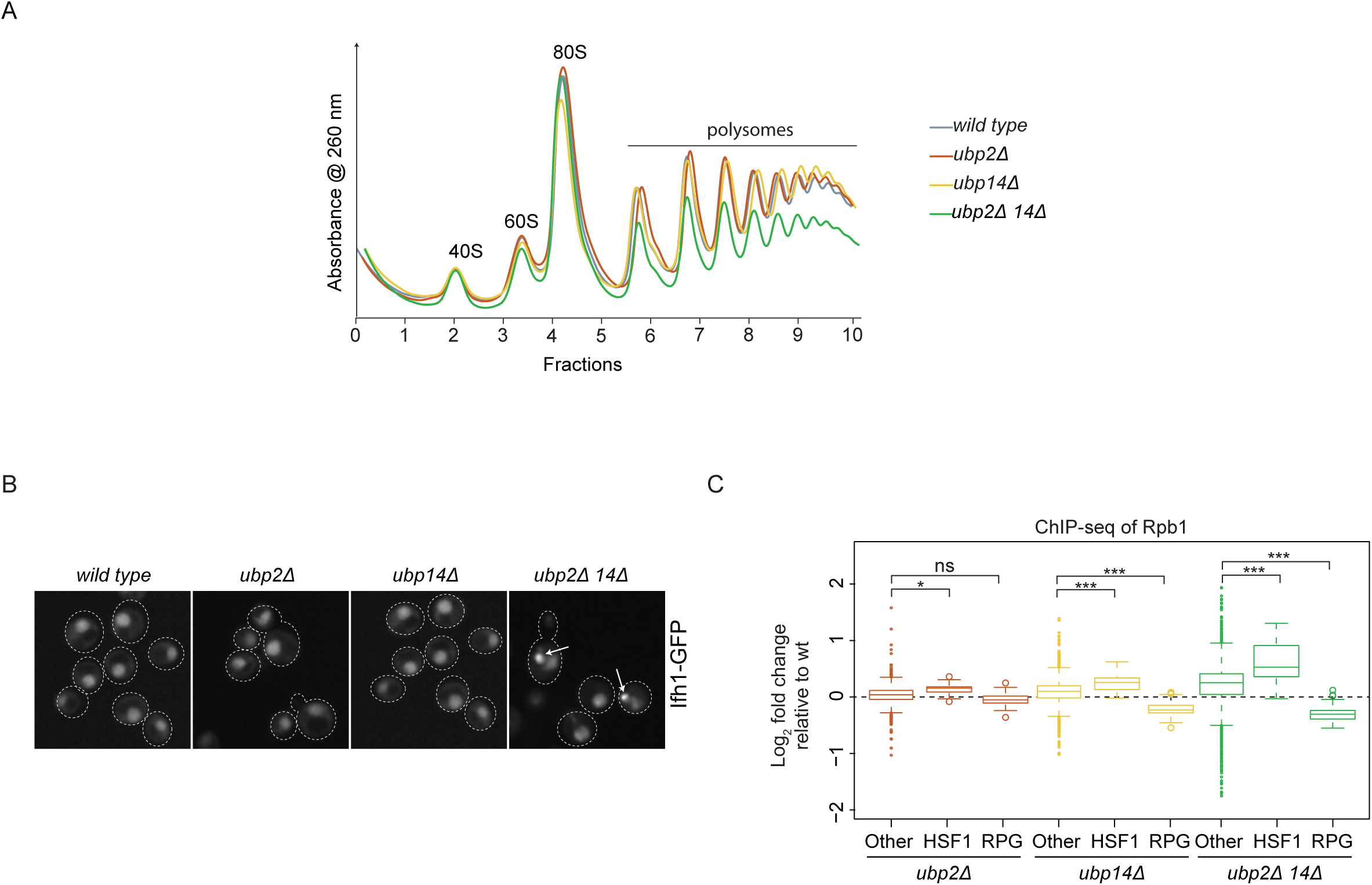
Ribosome biogenesis is disrupted in a *ubp2Δ ubp14Δ* double mutant. (A) Polysome sedimentation profiles (OD260) of wild type, *ubp2Δ*, *ubp14Δ* and *ubp2Δ ubp14Δ* strains are shown. Gradient fractions corresponding to 40S, 60S, 80S and polysomes are indicated. (B) Example micrographs of yeast cells expressing Ifh1-GFP in the indicated mutant strains. Arrows indicate Ifh1-GFP protein aggregates. (C) Box plots comparing RNAPII binding (as measured by Rpb1 ChIP-seq) in *ubp2Δ*, *ubp14Δ* and *ubp2Δ ubp14Δ* cells relative to *wild type* cells. Indicated gene categories include ribosomal proteins (RPG)-(n =139), Others (n = 4884) and HSF1 target genes (HSF1) (n = 19). ns = not significant; *= p < 0.05 and *** = p < 0.0001.

*</i>*Consistent with defects in ribosome biogenesis contributing to accumulation of ribosomal proteins at INQ in the *ubp2Δ ubp14Δ* double mutant, treating wild type cells with Diazaborine (DZA), a specific inhibitor of Drg1 which is an essential protein in 60S ribosomal subunit biogenesis, [69, 70], triggered all 36 INQ resident proteins (Figure 3A) to aggregate (representative examples shown in Figure S4C). Thus, a defect in ribosome biogenesis is a likely trigger for INQ formation in *ubp2Δ ubp14Δ* double mutant cells.

### *UBP2*- and *UBP14*-dependent protein aggregation and ribosome biogenesis defects depend on K29-linked unanchored polyUb chains

There are several documented examples where K48- and K63-linked unanchored polyubiquitin chains affect cell signaling processes by physically interacting with their effector proteins [36]. We used anti-Ub western blotting of polysome gradient fractions to show that K29-linked unanchored chains may similarly associate with downstream effectors. Specifically, a fraction of K29-linked unanchored polyUb chains co-sedimented with the 60S, 80S and polysome fractions, especially in cell extracts derived from a *ubp2Δ ubp14Δ* double mutant (Figure S5A). Moreover, expression of a Ub mutant defective in K29-polyUb formation (UbK29R) or deletion of the E3 ligase gene, *UFD4*, which leads to depletion of K29-linked unanchored polyUb chains (Figure 3), suppressed multiple protein homeostasis phenotypes that we observed in the *ubp2Δ ubp14Δ* double mutant cells: [1] the decreased levels of polysomes (Figure 5A, B); [2] Rpl23A-GFP and Ifh1- GFP aggregation (Figure 5C); [3] the rRNA processing defect (Figure 5D, Figure S5B), and [4] aggregation of all tested INQ substrates (Figure S5C). These findings suggest that free K29-linked unanchored polyUb chains that accumulate in the absence of *UBP2* and *UBP14*, negatively impact cell growth and ribosome biogenesis and these effects may be mediated via physical interactions with ribosomal proteins.

**Figure 5:**
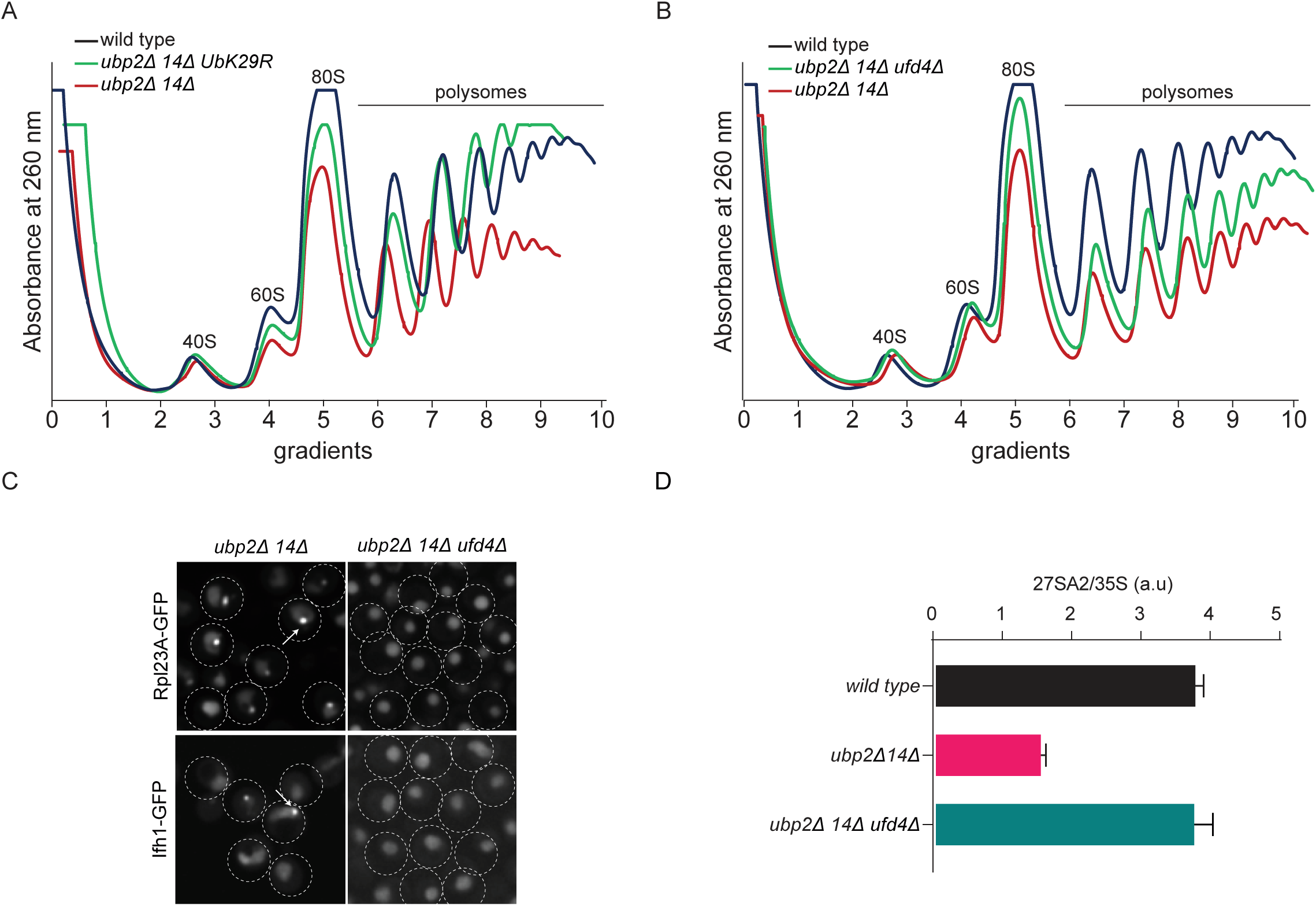
K29-linked free polyubiquitin chains co-sediment with ribosomes affecting their biogenesis. (A) Polysome sedimentation profiles (OD260) of wild type, *ubp2Δ ubp14Δ* and *ubp2Δ ubp14Δ* UbK29R strains are shown. In the latter strain, UbK29R is the sole source of Ub in the cell. Gradient fractions corresponding to 40S, 60S, 80S and polysomes are indicated. (B) Polysome profiles of wild type, *ubp2Δ ubp14Δ ufd2Δ* and *ubp2Δ ubp14Δ* strains are shown (labels as in (A)). (C) Example micrographs of yeast cells expressing Rpl23A-GFP or Ifh1-GFP in the indicated mutant strains. Arrows indicate GFP protein fusion aggregates. (D) Quantifications of the ratio of 27SA2 to 35S rRNA bands from a Northern blot to measure rRNA processing (arbitrary units; blot shown in Figure S5A). Error bars show standard deviation (n=2).

## Discussion

We used yeast functional genomics, proteomics and single cell imaging screens to reveal roles for the redundant DUBs *UBP2* and *UBP14* in preventing the accumulation of K29-linked unanchored polyUb chains, which causes toxicity by forming intracellular aggregates and by associating with ribosomes, disrupting their assembly and causing a severe growth defect. This disruption in ribosome biogenesis activates RASTR and promotes deposition of ribosomal proteins in the INQ compartment. Our study highlights a key role for K29-linked unanchored polyUb chains as a ‘barometer’ of proteostasis, a physiological role for INQ, and the redundant regulators that the cell deploys to regulate ribosome biogenesis (summarized in model in Figure 6).

**Figure 6:**
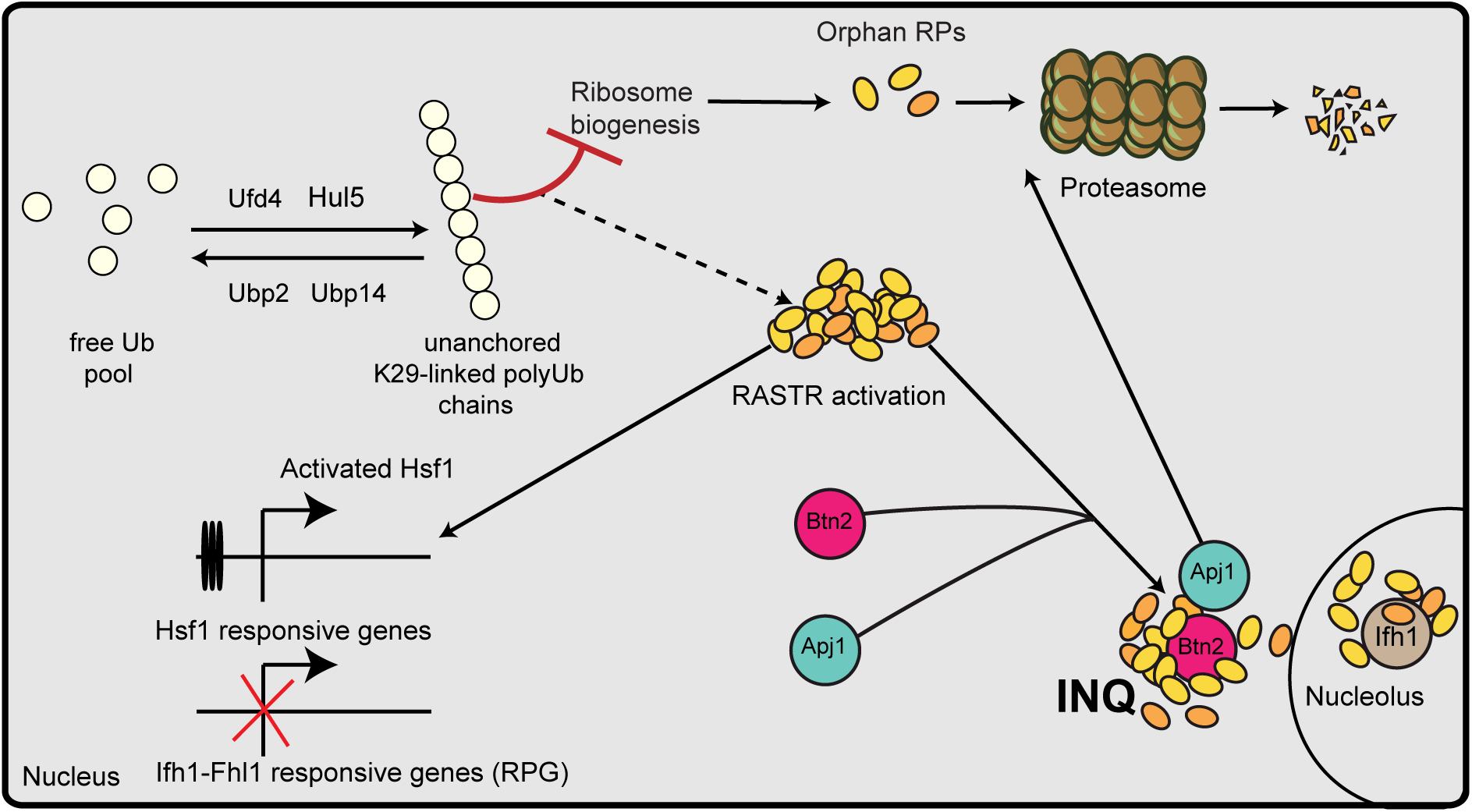
A model for ribosome biogenesis defects mediated by K29-linked polyubiquitin chains. Opposing the activity of Ubiquitin E3 ligases (Ufd4 and Hul5), DUBs Ubp2 and Ubp14 recycle the K29-linked unanchored polyUb chains and maintain the free Ub pool. In the absence of Ubp2 and Ubp14, K29-linked unanchored polyUb chains accumulate to form inclusions and disrupt ribosome biogenesis, which triggers the Ribosome assembly stress response (RASTR) that is characterized by activation Hsf1 target genes (such as *BNT2* and *APJ1*) and accumulation of numerous proteins, including ribosomal proteins, their assembly and associated factors, at the Intranuclear Quality control compartment (INQ). INQ accumulation occurs in a Btn2-dependent manner along with sequestration of the transcription factor Ifh1 at the nucleolus.

Our discovery that disruption of polyUb chain homeostasis by deleting *UBP2* and *UBP14* causes aggregation of K29-linked unanchored polyUb chains is consistent with studies in human cells showing that K29/K33-linked polyUb chains form intracellular inclusions [71]. Moreover, polyUb chains of various linkage types have been reported to have reduced thermostability, leading to aggregate formation *in vitro* and *in vivo* with possible relevance to human neurodegenerative diseases [49, 72]. However, under normal conditions, when their abundance is tightly regulated, K29-linked unanchored polyUb chains may facilitate ubiquitylation of toxic misfolded proteins targeted for degradation [36]. Consistent with a general proteostasis role, the abundance of unanchored polyUb chains increases under proteotoxic stress, which generates misfolded proteins [73] and polyUb chains of mixed topologies (i.e. K48 & K29 linked chains) are more potent in targeting proteins for degradation [74]. Interestingly, both Hul5 and Ufd4, which appear to be capable of synthesizing K29-linked unanchored polyUb chains *in vivo*, play a crucial role in degrading misfolded proteins under stress conditions [74–77].

Using a proteome-scale imaging screen, we discovered that the increased levels of K29-linked unanchored polyUb chains in the absence of *UBP2* and *UBP14* were associated with aggregation of ∼ 250 proteins (Table S3), the vast majority of which (201/259; ∼78%) had not been previously shown to form aggregates. These proteins were enriched for translation-related functions and included aggregates of several ribosomal proteins and their assembly partners in the nucleolus, nucleus and cytosol (Figures 3A, S4B). Such widespread protein aggregation likely reflects the combined effects of several defects associated with loss of *UBP2* and *UBP14*: (i) aberrant interactions of K29-polyUb chains and their aggregates with metastable proteins leading to protein misfolding; (ii) depletion of the free ubiquitin pool (Figure 1E); (iii) decreased ubiquitylation of proteins due to the depleted free ubiquitin pool (Figure 1C, Table S1); (iv) unappreciated K29- polyUb chain-dependent or other roles of *UBP2* and *UBP14*; (v) decreased turnover of polyubiquitylated substrates due to competition by abundant unanchored polyUb chains for binding to the proteasome [5]. Although, we did not observe any significant defect in the degradation of a proteasomal substrate, GFP-VHL (Figure S3B), substrate- and polyUb chain type-specific stabilization of proteins cannot be ruled out.

Regulation of ubiquitylation of the ribosomal protein Rps7 is key for efficient translation [32], and our proteomics study showed that Rps7 ubiquitylation was primarily observed in polysomes (Figure S5A) [78] and was enriched in the *ubp2Δ ubp14Δ* double mutant. Thus, the reduction in polysomes we observed in the *ubp2Δ ubp14Δ* may reflect the inefficient deubiquitylation of Rps7. However, *ubp14Δ* single mutants had no obvious change in polysome profiles yet had a similar level of enrichment for ubiquitylated Rps7, making it unlikely that increased ubiquitylation of Rps7 is responsible for the observed defects in polysome profiles and aggregation of proteins at INQ. However, a possible effect of Rps7 (and other RiBi factor) aggregation (Table S3) on the reduced polysomes that we observed in the *ubp2Δ ubp14Δ* double mutant cannot be ruled out.

As summarized in Figure 6, we discovered that increased levels of K29-linked polyUb chains impaired ribosome biogenesis by possibly physically associating with the pre-ribosomes, triggered RASTR and resulted in the aggregation of a subset of proteins (58/259; ∼22%) that were previously shown to be sequestered to INQ in response to DNA damage (MMS treatment) or proteotoxic stress[26-28, 79-81], suggesting that INQ formation is also induced in response to defective ribosome biogenesis. In addition to all previously known INQ substrates, we identified 10 novel substrates, including ribosomal proteins and their assembly factors, that localized to INQ in the absence of *UBP2* and *UBP14*, suggesting that these proteins may be targeted for proteasome-dependent degradation when ribosome assembly is impaired in the presence of unanchored K29 polyUb chains (Figure 6).

Several lines of evidence suggest that ribosome assembly stress may be a specific trigger of INQ formation. First, activation of RASTR or treatment with Diazaborine, which inhibits the essential Drg1 ATPase required for pre-60S ribosome maturation [69, 70], both lead to increased expression of the Btn2 chaperone required for INQ formation [24, 58] and aggregation of all INQ resident proteins reported in this study (representative examples shown in Figure S4C). Second, INQ formation is blocked upon Cycloheximide or Rapamycin treatment, which prevents protein synthesis and the accumulation of orphan ribosomal proteins [24, 28]. Finally, mutations in expansion segment 7 of the 25S rRNA gene lead to proteotoxic stress and cause a GFP-VHL reporter protein to form inclusions that also include rRNA and ribosomal proteins [82]. It is conceivable that when ribosome biogenesis is affected, a large number of highly abundant metastable ribosomal proteins and misfolded proteins accumulate due to errors in translation and simply overwhelm the protein homeostasis machinery, resulting in accumulation at the INQ compartment [19, 24, 82, 83]. However, why only a subset of specific proteins accumulate at INQ when ribosome biogenesis is impaired is not clear, and the mechanistic significance of their sequestration remains to be studied.

Previous studies demonstrated that polyUb chains with different linkage types can physically bind to substrate proteins leading to their aggregation and eventual clearance via autophagy [40, 49, 84, 85]. For example, K63-linked unanchored polyUb chains bind HDAC6 to activate aggregate clearance via autophagy [40, 84, 85]. Moreover, physical interactions between linear polyubiquitin chains and the NF-kB essential modulator, NEMO, facilitate liquid phase separation of NEMO, causing activation of IkB kinase (IKK), leading to NF-kB-mediated expression of many genes including those involved in immune and inflammatory responses [86]. Consistent with these studies, our data suggest that unanchored K29-polyubiquitin chains perturb ribosome biogenesis and/or assembly through direct physical interaction with ribosomes.

In human tissues, cells heterozygous for mutations in ribosomal genes exhibit a so-called ‘loser’ status and are eliminated by wild-type cells [87]. A RASTR-like response characterized by increased proteostasis stress may be the underlying mechanism behind loser status [25, 88] or apoptosis in cells line models of human diseases collectively termed “Ribosomopathies” [89]. These diseases are characterized by mutations in ribosomal genes that cause an imbalance in ribosomal protein stoichiometry and aggregation, characteristic features of the ribosome assembly defect that we observed in a *ubp2Δ ubp14Δ* double mutant. Conceptually similar negative interactions involving ribosomal mutations are seen in flies where growth defects associated with ribosomal mutations are enhanced by proteasome inhibition leading to cell death [88], and in yeast, where the global genetic network revealed negative genetic interactions connecting proteasome- and ribosome-encoding genes [44]. In addition, Rapamycin treatment alleviates the toxicity associated with Ribosomopathy mutations in cell lines [88, 89] and inhibition of TOR signaling prevents INQ formation in yeast [28]. Although the molecular mechanism underlying the cellular response to ribosomopathy-related mutations are not clearly understood, our findings highlight an interesting new link between ribosome biogenesis, INQ formation, protein aggregation and Ribosomopathies. Moreover, while mother cell retention of protein aggregates is a key mechanism of cellular ageing [60, 61, 90, 91], incomplete mother cell retention of protein aggregates observed in *ubp2Δ ubp14Δ* double mutant cells highlight a potential role for these DUBs and K29-linked unanchored polyUb chains in cellular ageing and rejuvenation.

## Acknowledgements

We thank Dr. Minglei Zhao for sharing the sAB-K29 antibody. This work was supported by grants from the National Institutes of Health (R01HG005853 to B.J.A., C.B.), and the Canadian Institutes of Health Research (PJT-180259 to B.J.A.). Equipment for automated image acquisition and analysis was purchased using funds from the Canadian Foundation for Innovation and the Ontario Research Fund. C.B. is a Fellow of the Canadian Institute for Advanced Research and B.J.A. holds a Tier 1 Canada Research Chair in Systems Genetics and Cell Biology.

## Author Contributions

H.G.S, M.C, C.B and B.J.A developed the concept for the paper and designed the bulk of the experiments. H.G.S performed the experiments, with the exception of those listed below, with help from E.S under the supervision of C.B and B.J.A. E.B performed the label free diglycine mass spectrometry-based proteomics experiments under the supervision of P.T. B.A performed the ChIP-seq experiments under the supervision of D.S. C.D and A.K.H performed polysome profiling, rRNA processing experiments and western blots of polysome profiling fractions.

M.P.D.M helped with organization and analysis of imaging data. C.P. aided in data analysis and representation. H.G.S, M.C, C.B and B.J.A wrote the paper with input from all the authors.

## Materials and Methods

### Yeast genetic techniques and strains Strain construction

*Saccharomyces cerevisiae* S288C strain and its derivatives (BY4741 or BY4742 or BY4743) were used in this study [92, 93]. Table S5 contains a complete list of strains including their genotypes. Most genetic manipulations including gene knockouts, N-terminal (promoter switching) and C-terminal epitope tagging (GFP) were carried out by homologous recombination with PCR-products encoding a selectable marker and carrying 45-50 bp homology on either end to the region of interest in the genome [94]. A list of all the primers used in this study are available in Table S5. Lithium acetate-polyethylene glycol-based transformation of recombinant DNA was employed in all cases [95].

For imaging screens, BY5299 (*MAT**α** can1pr::TDH3pr-tdTomato-NLS-URA3 lyp1pr::TDH3pr- E2Crimson-hphMX6 can1Δ::STE2pr-LEU2 his3Δ1 leu2Δ ura3Δ met15Δ lyp1Δ*) was used as the starting strain. E2Crimson and td-Tomato-NLS were used as cytosolic and nuclear markers respectively. The starting strain was used to construct the following query test strains: *MAT*α *ubp14Δ:natMX4 URA3pr::MAL63-kanMX4-MAL32pr* (single mutant)*, MAT*α *ubp2Δ::natMX4 URA3pr::MAL63-kanMX4-MAL32pr* (single mutant) and *MAT*α *ubp14Δ::natMX4 UBP2pr::Mal63-kanMX4-MAL32pr* (double mutant). Native promoters were switched to MAL32pr via PCR-mediated promoter switching using pMaM458 as template [51].

### Yeast growth assay in liquid medium

Yeast cultures were grown overnight in SC-Maltose (1.7 g/L yeast nitrogen base without amino acids and ammonium sulphate, 1 g/L monosodium glutamic acid, 20 g/L Maltose, amino acid dropout mix). Cultures were diluted 200 times in fresh media (either SC-Maltose or SC-Dextrose) in a 96 well plate and growth was monitored every 30 min for 24 hrs by measuring the absorbance at 600 nm in an automated shaker and plate reader set-up (S&P robotics). Experiments were performed in three biological replicates.

### Yeast spot titer assay

Yeast cultures were grown overnight to saturation in minimal media (1.7 g/L yeast nitrogen base without amino acids and ammonium sulphate, 1 g/L monosodium glutamic acid, 20 g/L Maltose, amino acid dropout mix). 200 μl of each culture corresponding to OD600 = 1.0 were transferred to a 96 well plate, serially diluted (1:5 dilutions) with water and subsequently pinned onto agar plates using a spotter.

### High-throughput microscopy

Yeast cultures were prepared for microscopy and imaged as previously described [21, 52, 97]. Briefly, the *MAT***a** wild type and mutant query strains described above expressing ORF-GFP fusion proteins (from yeast GFP collection [55]) were grown at 30°C and imaged in low fluorescence synthetic minimal medium supplemented with appropriate drugs and 2% glucose. Cells were transferred to 384-well PerkinElmer CellCarrier Ultra imaging plates and centrifuged for 1 minute at 500 g before imaging. Micrographs were obtained on an Opera Phenix (PerkinElmer) automated confocal microscope. All imaging was done with a 60× water-immersion objective. GFP was excited using a 488 nm laser and emission collected through a 520/35-nm filter. tdTomato was excited using a 561-nm laser, and emission collected through a 600/40-nm filter. E2crimson was excited using a 640 nm laser, and emission collected through a 690/50 nm filter. Images were analyzed by eye to identify cellular factors that aggregate in the DUB mutants.

### Immunofluorescence microscopy

Yeast cells were grown overnight to saturation in synthetic dextrose medium (1.7 g/L yeast nitrogen base without amino acids and ammonium sulphate, 1 g/L monosodium glutamic acid, 20 g/L Glucose, amino acid dropout mix). Cultures were diluted to an OD600 = 0.1 in fresh medium (50 ml) and grown until an OD600= 0.8. Formaldehyde solution (0.5 ml of 37% (v/v)) was added to the cultures and incubated for 15 min on a shaker. Cells were spun at 2000 rpm for 5 min and resuspended in 10 ml of 4% (w/v) Paraformaldehyde solution made with 100 mM KPO4 buffer pH 6.5 and incubated for 1 hour at room temperature with intermittent mixing. Cells were harvested and washed with digestion buffer (1.2 M sorbitol in 100 mM KP04 buffer, pH 6.5). Cells were resuspended in 1 ml of digestion buffer, and 5 µl of β-mercaptoethanol and 10 µl Zymolase T-100 (from 10 mg/ml stock) were added to the resuspended cells and incubated for 30 min at 30°C.

Spheroplasts were harvested by centrifugation at 2000 rpm for 5 min and resuspended in 1 ml of 100 mM KPO4 buffer (pH 6.5) and placed on ice. 10 µl of resuspended cells were added to 0.1 % (w/v) poly-lysine coated slides and incubated for 10 min. Unbound cells were aspirated and washed a few times with blocking buffer (100 mM KPO4 buffer pH 6.5, containing 5% (w/v) Bovine Serum Albumin). Spheroplasts were incubated with blocking solution for 1 hour and then washed 3 times with blocking buffer containing 1% (v/v) Triton X-100 and 3 times with just the blocking solution. Spheroplasts were treated with blocking solution containing sAB-K29 [50] for 2 hrs (at a final concentration of 1 µg/ml; a kind gift from Minglei Zhao, University of Chicago), washed a few times with blocking solution and treated with secondary antibody (Alexa Fluor® 647 AffiniPure Goat Anti-Human IgG, F(ab’)2 fragment specific. Cat# AB_2337881) for 1 hr. The spheroplasts were washed 5 times in KPO4 buffer and mounted using ProLong™ Diamond

Antifade Mountant with DAPI (Thermofischer; cat# P36971) before imaging. Images were acquired with Opera Phenix (PerkinElmer) automated confocal microscope. All imaging was done with a 60× water-immersion objective. Samples were excited using a 640 nm laser, and emission collected through a 690/50 nm filter.

### Isolation of ubiquitylated proteins using Tandem Ubiquitin Binding Entities (TUBEs)

Logarithmically growing cells in minimal media (1 L) were harvested, washed twice with cold distilled water and finally resuspended in cold lysis buffer (50 mM Tris-HCl pH 8.0,150 mM NaCl, 0.2 mM EDTA, 5% (v/v) glycerol, 1 mM DTT, 0.2 % Tween-20, 5 mM chloroacetamide, 10 mM N-ethyl malemide, 2X Halt protease inhibitor cocktail (Thermofischer; Cat# 78429)). Acid washed glass beads (Sigma; Cat# G8772) were added and cells lysed by bead-beating for 10 min at a shaking speed of 3450 oscillations/min using a BioSpec Mini-Beadbeater with intermittent cooling of samples on ice. Lysate was then clarified by first centrifuging at 3000 g for 5 min and subsequently at 16000 g for 15 min. 120 ul of Agarose-TUBE2 (Cat# UM402; lifesensors) was added to cell lysate corresponding to 10 μg of protein (as measured by Bradford’s method) and incubated on a rotater for 90 min at 4°C. The agarose beads were washed four times with lysis buffer and finally the bound protein was eluted by heating (95 °C for 5 min) in Laemmli sample loading buffer. Anti-Zwf1 antibodies for the loading control were purchased from Sigma (Cat# A9521).

### K-ε-GG (di glycine) antibody-based isolation of ubiquitylated peptides and their label free quantification using mass spectrometry

Yeast strains were individually grown in 1 L of SC-Dextrose medium to OD600=0.8 at 30°C in a shaking incubator. Cells were harvested by centrifugation at 4000 g for 5 min and washed with ice cold water. Wet weight of cell pellets was measured. For every 1g of cell pellet, 3g of ice-cold glass beads (425-600 um; sigma; Cat# G8772) and 2 ml of lysis buffer (100 mM HEPES pH 7.5, 150 mM NaCl, 0.2 mM EDTA, 1 mM DTT, 10% (v/v) glycerol, 2X Halt^TM^ Protease Inhibitor Cocktail (Thermofischer; Cat# 78438), 10 mM 2-chloroacetamide) were added and cells were lysed by vortexing for 12 min (with 1 min on ice after every 1 min of vortexing). NP40 detergent was added to a final concentration of 0.2% (v/v) and the lysate was incubated on a rotator for 10 min at 4°C. Unlysed cells were removed by centrifugation at 3000 g for 5 min. Lysates were further clarified by two rounds of centrifugation at 17500 rpm for 10 min each. Protein concentrations were measured by Bradford’s method. Lysates were then TCA precipitated, washed twice with ice-cold acetone, and air-dried at 37°C for 30 min. Protein pellets were resuspended in lysis buffer (Tris 50 mM Urea 8M, TCEP 5mM and chloroacetamide 55 mM). Samples were sonicated and vortexed for 1 hour. Protein quantitation was performed by Bradford assay. For each sample, 10 mg of protein were digested with trypsin (enzyme/total protein: 1/25) overnight at 37°C with mild vortexing. Samples were desalted on Oasis HLB columns (Waters, Cambridge, MA) and dried down.

The PTMScan ubiquitin branch motif (K-ε-GG) immunoaffinity beads (Cell Signaling Technology, Danvers, MA) from one vial were washed three times with 1 mL of 200 mM triethanolamine pH 8.3. For cross-linking, 1 mL of 20 mM dimethyl pimelimidate (DMP) in 200 mM triethanolamine pH 8.3 was added and the sample was incubated for 1 h at room temperature under gentle rotation. The reaction was quenched by adding 50 μL of 1 M Tris–HCl pH 7.5 and incubating for 30 min under gentle rotation. Cross-linked beads were washed three times with ice-cold PBS, resuspended in 300 μL of PBS, and stored at 4 °C. For the ubiquitin IP, digests were reconstituted in 1 mL of PBS and incubated with 100 μL of anti-K(GG)-coupled agarose beads in PBS for 2 h at 4 °C under gentle rotation. Anti-K(GG)-coupled beads were washed two times with 1 mL of IAP buffer (50 mM 3-(*N*-morpholino) propanesulfonic acid (MOPS)/NaOH, 10 mM Na2HPO4, 50 mM NaCl, pH 7.2) and three times with 1 mL of ice-cold water. Ubiquitylated peptides were eluted two times with 55 μL of 0.2% formic acid in water and filtered through a 0.45 μm spin tube. Eluted peptides were dried down by SpeedVac and stored at −80 °C for MS analysis.

Peptides were loaded and separated on a home-made reversed-phase column (150-μm i.d. by 200 mm) with a 56-min gradient from 10 to 30% ACN-0.2% FA and a 600-nl/min flow rate on a Easy nLC-1000 connected to an Orbitrap Fusion (Thermo Fisher Scientific, San Jose, CA). Each full MS spectrum acquired at a resolution of 60,000 was followed by tandem-MS (MS-MS) spectra acquisition on the most abundant multiply charged precursor ions for a maximum of 3s. Tandem-MS experiments were performed using collision-induced dissociation (CID) at a collision energy of 30%. The data were processed using PEAKS X (Bioinformatics Solutions, Waterloo, ON) and a Uniprot human database (20349 entries). Mass tolerances on precursor and fragment ions were 10 ppm and 0.3 Da, respectively. Fixed modification was carbamidomethyl (C). Variable selected posttranslational modifications were oxidation (M), deamidation (NQ), phosphorylation (STY) and GlyGly(K). LOESS normalization was achieved with the R package Normalizer [96].

### Cycloheximide chase experiments and western blotting

Cells were grown to logarithmic phase and an aliquot of cells corresponding to OD600 =3 were harvested, washed in cold distilled water and frozen in liquid nitrogen. Cycloheximide was added to the culture to a final concentration of 100 μg/ml. Cultures were then incubated on a shaker and aliquots of cells were taken at different time points. Frozen cells were resuspended in 1 ml of cold distilled water, then 185 μl of 1.85M NaOH and 10 μl of β-mercaptoethanol were added and the mixture incubated on ice for 10 min. 150 μl of 50 % (w/v) trichloroacetic acid were added followed by further incubation on ice for 10 min. Samples were then centrifuged at 16000 g for 15 min and the pellets were resuspended in sample loading buffer containing 8M urea and heated at 65 ^0^C for 10 min. Samples were run on Biorad 4-20% gradient gels and transferred to PVDF membranes using a Biorad Trans-blot system. Membranes were blocked in 5 % (w/v) milk powder and probed with the relevant primary antibodies: P4D1 horseradish peroxidase (HRP) conjugate anti-Ub antibody was used (Santa cruz; Cat # Sc-8017 HRP) at 1:1000 dilution, anti-Ub antibody (cell signaling; cat# 3933), living colours GFP monoclonal antibody (Takara, Cat# 632375) at 1:5000 dilution and anti-ZWF1 antibody (sigma; Cat# A9521) at 1:10000 dilution. The corresponding rat anti-mouse (Biorad; Cat# 1706516) or goat anti-rabbit (Biorad; Cat# 1706515) secondary antibodies (IgG(H+L)-HRP) were used at 1:10000 dilution for detection using a Versadoc system (Biorad).

### Polysome profiling

Cycloheximide (Sigma; Cat# 0180) was added directly to the exponentially growing yeast culture medium to a final concentration of 50 µg/mL. Cells were collected by centrifugation, rinsed with buffer K [20 mM Tris-HCl pH 7.4, 50 mM KCl, 10 mM MgCl2] supplemented with 50 µg/mL cycloheximide and collected again by centrifugation. Dry pellets were resuspended with approximately one volume of ice-cold buffer K supplemented with 1 mM DTT, 1 × Complete EDTA-free protease inhibitor cocktail (Roche; Cat# 04693132001), 0.1 U/µL RNasin (Promega; Cat# N2111) and 50 µg/mL cycloheximide. About 250 µL of ice-cold glass beads (Sigma; Cat# G8772) were added to 500 µL aliquots of the resuspended cells and cells were broken by vigorous shaking, three times 2 min, separated by 2 min incubations on ice. Extracts were clarified through two successive centrifugations at 13,000 rpm and 4°C for 5 min and quantified by measuring absorbance at 260 nm. About 30 A260 units were loaded onto 10–50% sucrose gradients in buffer K, and then centrifuged for 150 min at 39,000 rpm and 4°C in an Optima L-100XP Ultracentrifuge (Beckman-Coulter) using a SW41Ti rotor. Following centrifugation, 18 fractions of 500 µl each were collected from the top of the gradients with a Foxy Jr. apparatus (Teledyne ISCO). The absorbance at 254 nm was measured during collection with a UA-6 device (Teledyne ISCO).

### Protein extractions from sucrose gradient fractions

Proteins contained in sucrose gradient fractions were extracted as follows: 36 µl of 100 % (w/v) trichloroacetic acid were added to 200 µL of each crude sucrose gradient fraction (final concentration 15%) along with 1 µl 20 mg/ml glycogen, mixed thoroughly and incubated for 1 hour on ice. Samples were centrifuged at 13,000 for 20 min at 4°C. Protein pellets were rinsed with 500 µl acetone and centrifuged at 13,000 rpm for 5 min at 4°C. Protein pellets were air dried and resuspended with 50 µl SDS loading buffer.

### Western blotting of polysome fractions

Protein samples were run through 10% SDS-Polyacrylamide gels and transferred to nitrocellulose membranes using a Trans-Blot Turbo apparatus (BioRad). Membranes were saturated for 1 h with PBST buffer (137 mM NaCl, 2.7 mM KCl, 10 mM Na2HPO4, 2 mM KH2PO4, 0.1% Tween-20) containing 5% (w/v) powder milk. Following incubation for 2 h with the same buffer containing the primary antibodies, membranes were rinsed 3 times for 5 min with PBST buffer, incubated for 1 h with the secondary antibodies diluted in PBST containing 5% (w/v) powder milk and finally washed three times for 10 min with PBST buffer. Luminescent signals were generated using the Clarity Western ECL Substrate (Bio-Rad) and captured using a ChemiDoc imager (Bio-Rad).

### Analysis of pre-rRNA processing by Northern blotting

Extractions of total RNA from yeast cells were performed as described previously[97]. Equal amounts of total RNA (4 µg) were run through agarose gels as described [98]. RNAs were transferred to Amersham Hybond N^+^ membranes (GE Healthcare) by capillarity and membranes were hybridized with a ^32^P-labelled oligonucleotide probe (23S.1, 5’- GATTGCTCGAATGCCCAAAG-3’) using Rapid-hyb buffer (GE Healthcare). Radioactive membranes were exposed to Phosphorimaging screens and analysed using a Typhoon Tryo (GE Healthcare).

### ChIP-Seq

Yeast cultures of 50 mL in complete medium were collected at OD600 = 0.4–0.6 for each condition. The cells were crosslinked with 1% formaldehyde for 10 min and quenched by adding 125 mM glycine for 5 min at room temperature. Cells were then washed with ice-cold HBS (50 mM HEPES-Na pH:7.5, 140 mM NaCl) and resuspended in 0.6 ml of ChIP lysis buffer (50 mM HEPES-Na pH:7.5, 140 mM NaCl, 1 mM EDTA, 1% NP-40, 0.1% sodium deoxycholate) supplemented with 1 mM PMSF and 1× protease inhibitor cocktail (Roche; Cat# 04693132001). The cells were broken using Zirconia/Silica beads (BioSpec) and lysates were centrifuged at 13,000 rpm for 30 min at 4°C. Following centrifugation, the pellets were resuspended in 300 μl ChIP lysis buffer containing 1 mM PMSF and sonicated for 15 min (30s ON–60s OFF) in a Bioruptor (Diagenode). The lysates were centrifuged at 7000 rpm for 15 min at 4°C, following which primary antibodies were added to the supernatant and incubated for 1 h at 4°C. ChIP was performed using Abcam ab5131 for RNAPII. Magnetic beads coupled to IgG against rabbit (Dynabeads™ M-280 Sheep Anti-Rabbit; Thermofischer; Cat# 11203D) were washed three times with PBS (137 mM NaCl, 2.7 mM KCl, 10 mM Na2HPO4, 1.8 mM KH2PO4) containing 0.5% BSA and added to the lysates (30 μl of beads/300 μl of cell lysate). The samples were incubated for 2 h at 4°C. The beads were washed twice with AT1 buffer (50 mM HEPES-Na pH: 7.5, 140 mM NaCl, 1 mM EDTA, 0.03% SDS), once with AT2 buffer (50 mM HEPES-Na pH: 7.5, 1 M NaCl, 1 mM EDTA), once with AT3 buffer (20 mM Tris–Cl pH: 7.5, 250 mM LiCl, 1 mM EDTA, 0.5% NP-40, 0.5% sodium deoxycholate) and twice with TE buffer (10 mM Tris, 1 mM EDTA). The chromatin was eluted from the beads by resuspension in TE containing 1% SDS and incubation at 65°C for 10 min. The eluate was transferred to new Eppendorf tubes and incubated overnight at 65°C to reverse the crosslinks. The DNAs were purified using the MinElute PCR Purification Kit (Qiagen; Cat# 28004). DNA libraries were prepared using TruSeq ChIP Sample Preparation Kit (Illumina) according to manufacturer’s specifications. The libraries were sequenced using an Illumina HiSeq 2500 platform at the Institute of Genetics and Genomics of Geneva (iGE3; http://www.ige3.unige.ch/genomics-platform.php) and the reads were mapped to the sacCer3 genome assembly using HTSStation60 (read densities were calculated using shift: 100, extension: 50 bp). All densities were normalized to 10M reads. The signal was quantified for each gene between the transcription start site (TSS) and transcription termination site (TTS). All data from publicly available databases were mapped using HTS Station (http://htsstation.epfl.ch; [99]). Values of ChIP-seq signal for each gene are reported in Table S4.

### GO slim biological process

We performed the GO biological process enrichment analysis for each set of SPMs using the GO Slim mapping file available through the Saccharomyces Genome Database (www.yeastgenome.org/). SciPy’s hypergeometric discrete distribution package was used to calculate P-values. P-values were adjusted using the Bonferroni correction. Fold enrichment was calculated as (mutants in term/all mutants)/(term size/all background).

---

**Figure S1:**
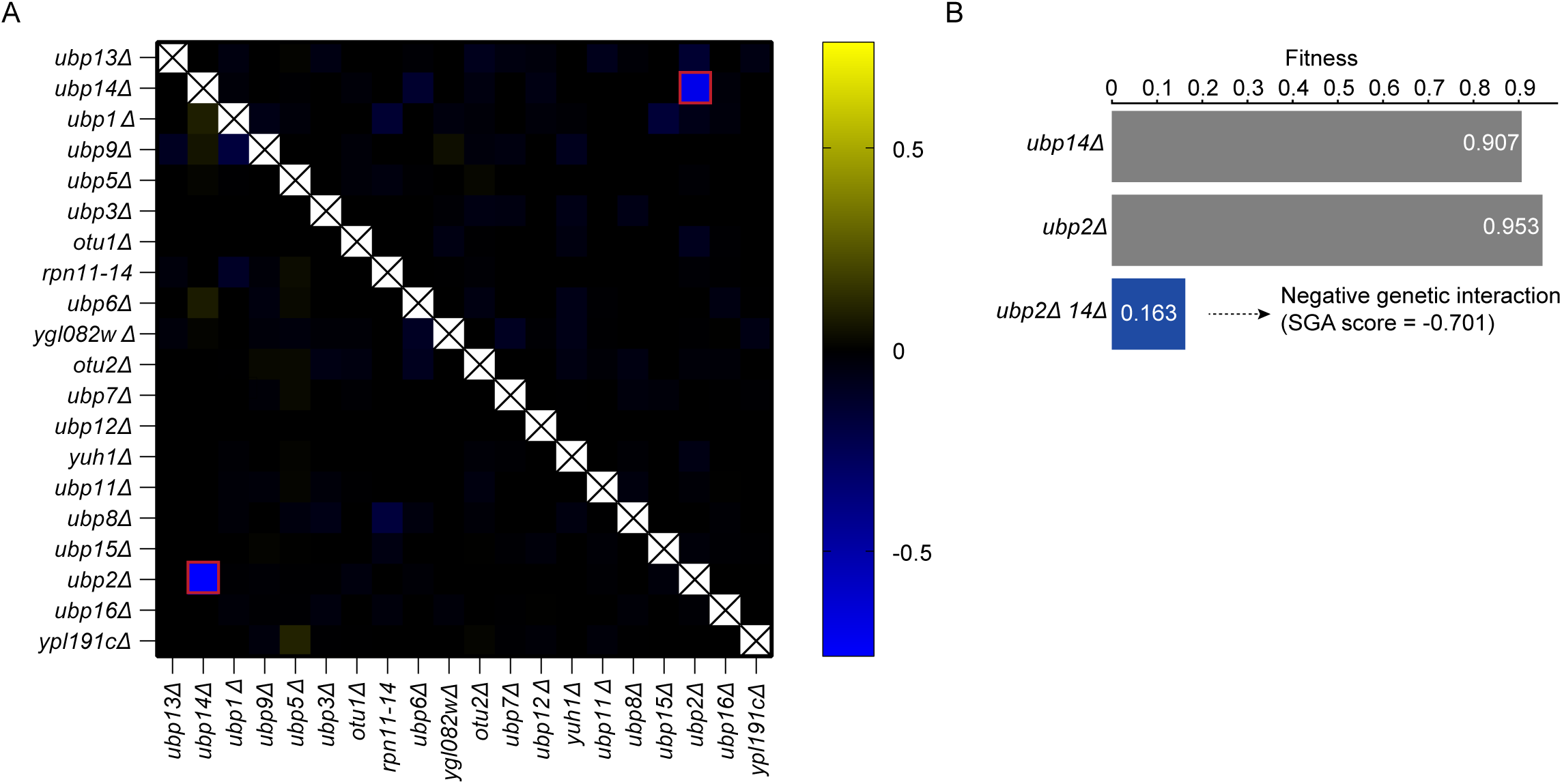
Genetic redundancy of DUBs. (A) A matrix of genetic interaction scores for all tested DUB gene pairs derived from the global genetic interaction network [44, 45]. Negative genetic interactions are coloured blue and positive interactions are coloured yellow (the scale bar shows the genetic interaction score (SGA score) [44]). A negative interaction in the *ubp2Δ ubp14Δ* double mutant is marked by a red box. (B) Fitness of *ubp2Δ, ubp14Δ* and *ubp2Δ ubp14Δ* mutants based on colony size measurements. The numbers in the bars indicate the colony size measurement for each mutant relative to wild-type [44]. The fitness of the *ubp2Δ ubp14Δ* double mutant is shown in the blue bar and is much less than expected based on the product of the single mutant fitness measurements, indicated a negative genetic interaction [44, 45].

**Figure S2:**
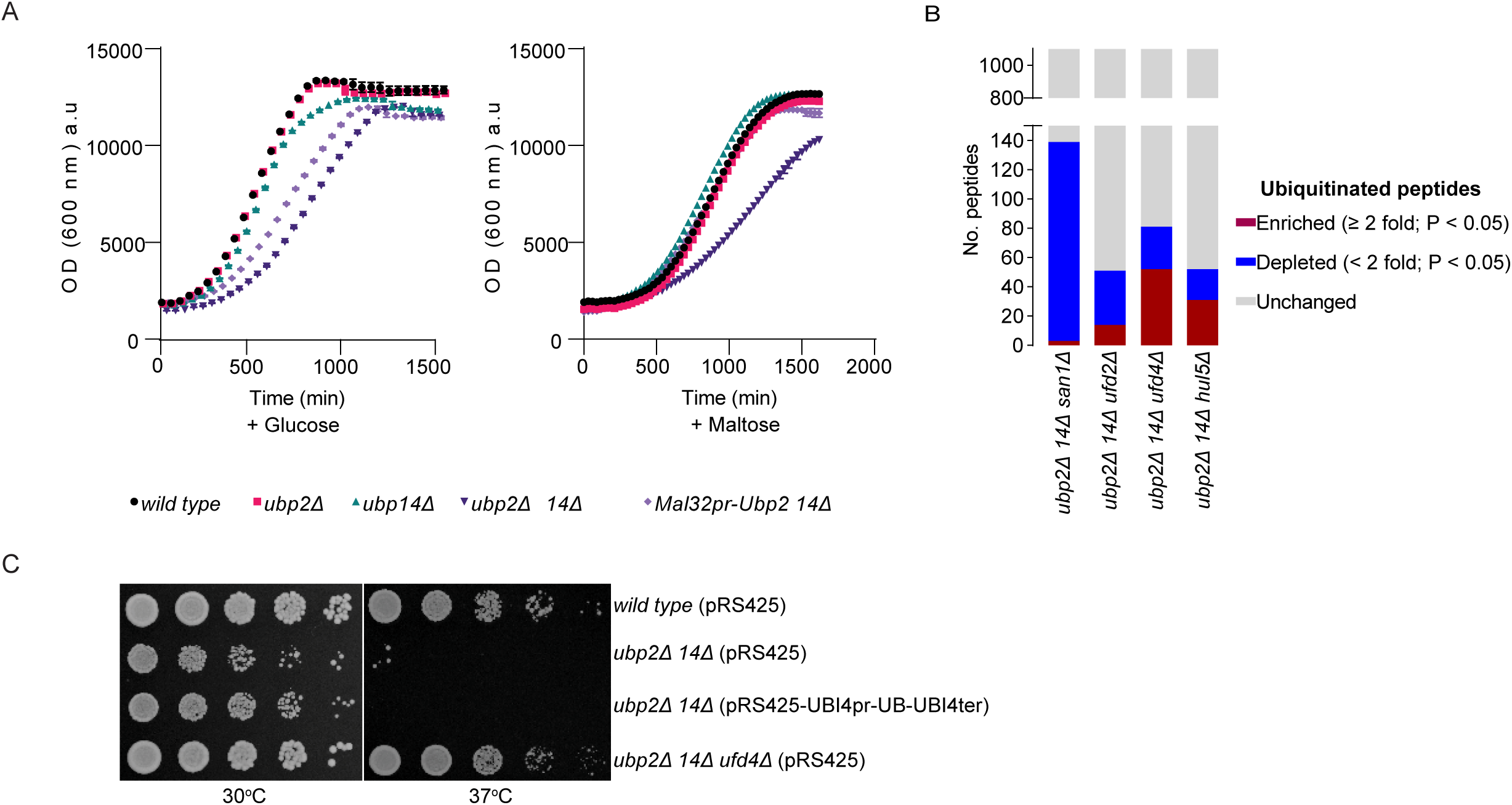
DUB-E3 ligase that function with Ubp2 and Ubp14 to regulate K29-linked unanchored poly Ub chains. (A) Liquid growth assays of *ubp2Δ, ubp14Δ* and *ubp2Δ ubp14Δ* mutants expressing a glucose-repressible and maltose-inducible allele of *UBP*2 (*MAL32*pr-*UBP2).* The indicated strains (legend at bottom) were diluted into fresh medium containing either glucose or maltose and growth was monitored every 30 minutes by measuring the optical density (OD) at an absorbance of 600 nm. Experiments were performed in biological triplicate and the average value is shown with error bars indicating the standard deviation. (B) Label-free quantitative mass spectrometry on ubiquitylated peptides isolated using Lys-ℇ-Gly- Gly (diGly) antibodies (n=3). Fraction of peptides that are enriched (≥2-fold, P < 0.05; blue) or depleted (<2-fold, P < 0.05; red) for ubiquitin modification in *ubp2Δ ubp14Δ san1Δ*, *ubp2Δ ubp14Δ ufd2Δ, ubp2Δ ubp14Δ ufd4Δ and ubp2Δ ubp14Δ hul5Δ* cells relative to *ubp2Δ ubp14Δ* cells are shown. The fraction of proteins whose ubiquitination status remained unchanged in the indicated mutant relative to the *ubp2Δ ubp14Δ* mutant is shown in grey. (C) Serial spot dilutions of wild-type or *ubp2Δ ubp14Δ* mutant strains transformed with empty vector (pRS425) or a ubiquitin-overexpressing plasmid (pRS425-*UBI4*pr-*UB*-*UBI4*ter). Plates were incubated at 30°C or 37°C as indicated.

**Figure S3:**
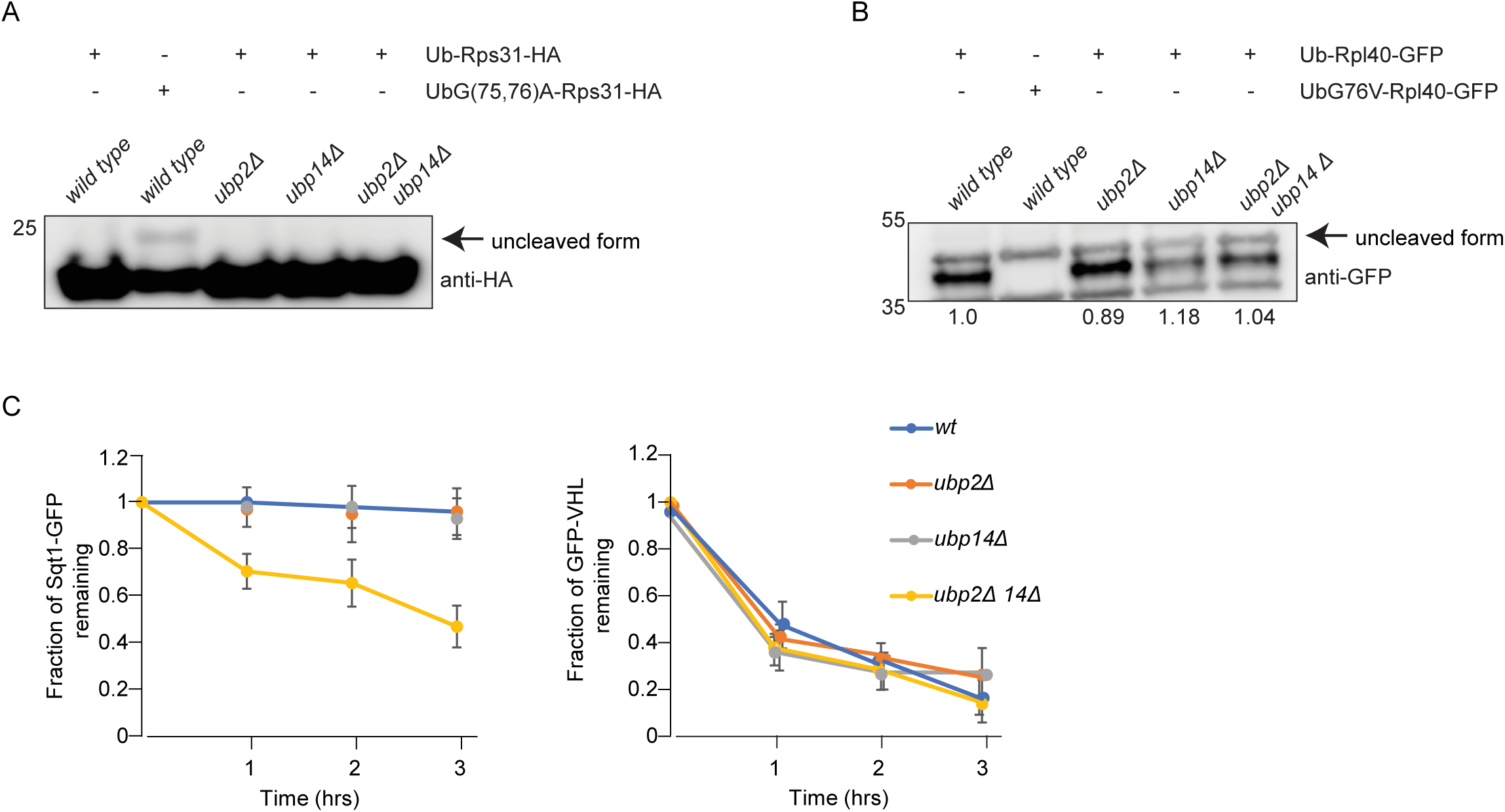
Protein aggregation is not explained by improper cleavage of Ub-ribosomal fusion proteins or a general defect in protein degradation. (A) anti-HA immunoblot of protein extracts derived from the indicated strains expressing Ub- Rps31-HA or a corresponding ubiquitin cleavage defective variant (UbG(75,76)A-Rps31-HA). Accumulation of an uncleaved version of Ub-Rps31 was only apparent when the uncleavable variant was expressed in wild-type cells and is indicated with the arrow. (B) Anti-GFP immunoblot of protein extracts from the indicated strains expressing Ub-Rpl40-GFP or a corresponding ubiquitin cleavage defective version (UbG76V-Rpl40-GFP). The uncleaved form of Ub-Rpl40 is indicated with an arrow. (C) Cycloheximide chase-based monitoring of degradation kinetics of Sqt1-GFP and GFP-VHL (an INQ substrate) in the indicated DUB mutants. Protein levels were assessed based on quantification of anti-GFP immunoblotting. Error bars indicate standard deviations (n=3).

**Figure S4:**
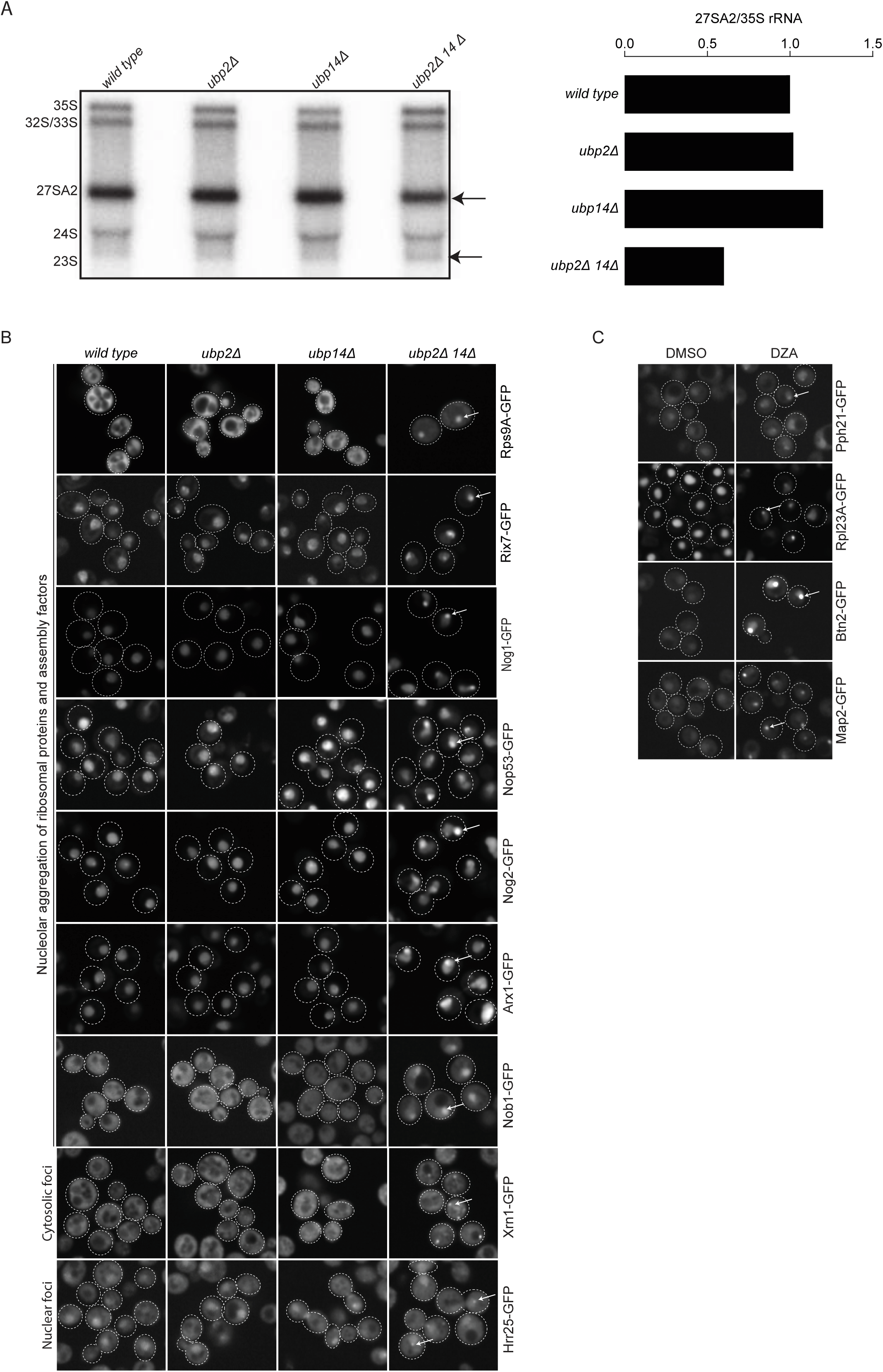
*ubp2Δ ubp14Δ* cells are defective in ribosome biogenesis. (A) Northern blot to monitor rRNA processing in the indicated strains (left panel). A decrease in 27SA2 rRNA (indicated by arrow) and concomitant increase in 23S rRNA relative to 35S rRNA in the *ubp2Δ ubp14Δ* double mutant indicates an rRNA processing defect. The graph on the right shows quantification of the relative signal of 27SA2/35S RNA on the Northern blot. (B) Example micrographs of yeast cells expressing specific GFP-fusion proteins (listed to the right of the micrographs) in the indicated mutant strains. Arrows highlight GFP-protein aggregates. (C) Example micrographs of yeast cells expressing specific GFP-fusion proteins in either the presence of a vehicle (DMSO) or Diazaborine (DZA), a specific inhibitor of Drg1, which is essential for 60S ribosome biogenesis. Arrows indicate GFP-protein aggregates. DZA was used at a final concentration of 7.5 µg/ml and cells were incubated for 2 hrs with shaking at 30°C before images were acquired using a confocal microscope.

**Figure S5:**
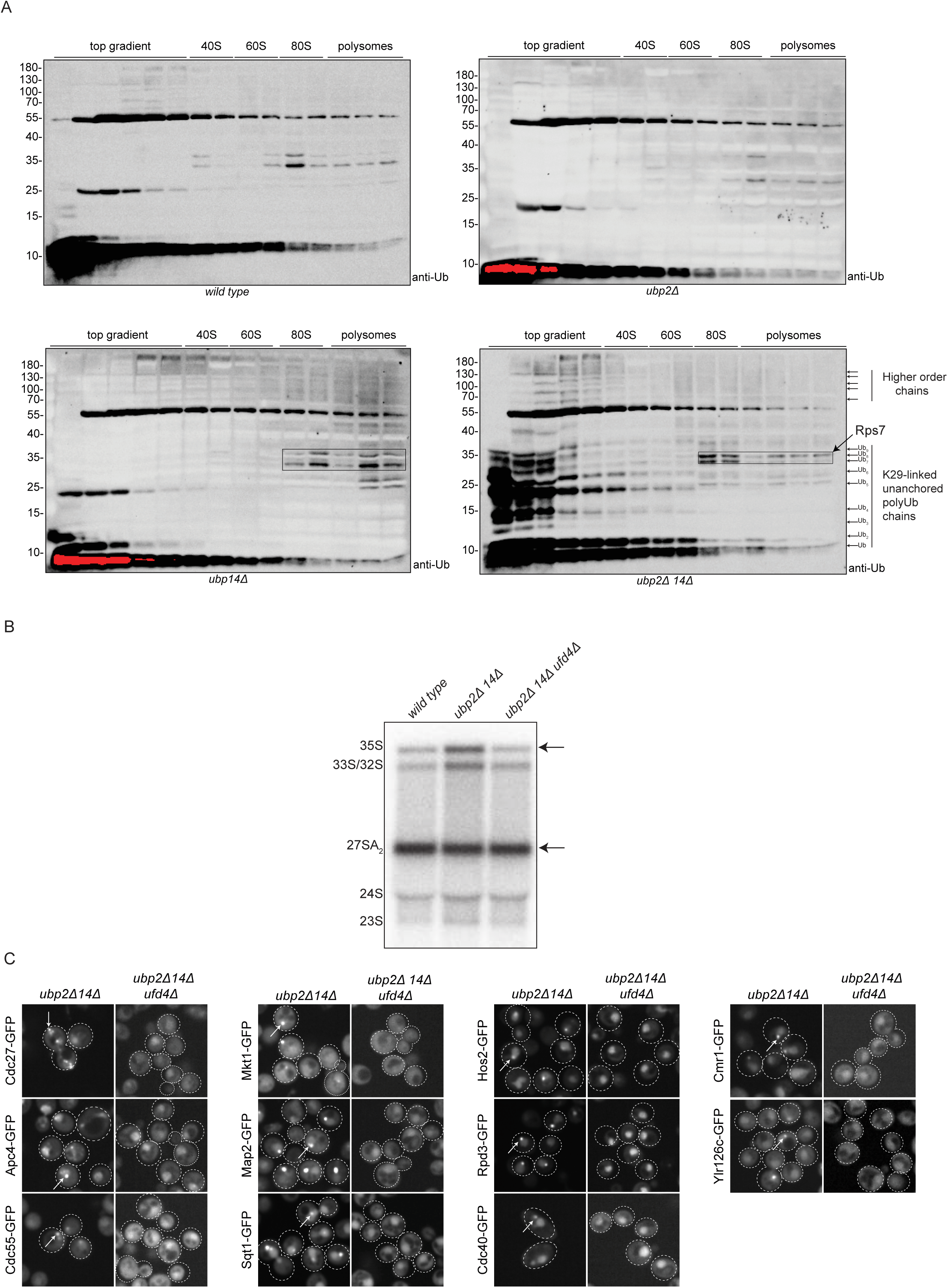
Loss of *UFD4* suppresses INQ formation. (A) Anti-Ub antibody immunoblot analysis of polysome fractions derived from a *wild type* strain, as well as *ubp2Δ, ubp14Δ* and *ubp2Δ ubp14Δ* mutant strains. Position of the blotted fraction on the polysome gradient is shown at the top of the blot, while the identity of various Ub derivatives is indicated to the right of the *ubp2Δ ubp14Δ* blot. The position of molecular weight markers is shown to the left of each blot. Protein bands within the boxed area in the 80S and polysome fractions likely represent ubiquitylated forms of Rps7A/B. (B) Northern blot to monitor rRNA processing in the indicated strains. The 35S rRNA and 27SA2 rRNA bands are highlighted with arrows, and their ratio in the different strains is quantified in Figure 5D. (C) Loss of *UFD4* suppresses aggregation of numerous INQ substrates in a *ubp2Δ ubp14Δ* double mutant. The tested INQ-GFP substrates are indicated to the left of the micrographs and the relevant genotype (*ubp2Δ ubp14Δ* or *ubp2Δ ubp14Δ ufd4*Δ) is shown at the top of the figure. Inclusions are highlighted with arrows.

**Movie S1:** Time-lapse movie monitoring GFP-VHL inclusions at 15 min intervals in logarithmically growing *ubp2Δ* cells.

**Movie S2:** Time-lapse movie monitoring GFP-VHL inclusions at 15 min intervals in logarithmically growing *ubp14Δ* cells.

**Movie S3:** Time-lapse movie monitoring GFP-VHL inclusions at 15 min intervals in logarithmically growing *ubp2Δ ubp14Δ* cells.

Supplementary tables and movies can be accessed at the below link: https://1drv.ms/f/s!AinYpOwRdgjOgldyzmsqNXUdRvqB?e=u4XtKF

## References

1. Rape, M., Ubiquitylation at the crossroads of development and disease. Nat Rev Mol Cell Biol, 2018. 19(1): p. 59–70.

2. Komander, D. and M. Rape, The ubiquitin code. Annu Rev Biochem, 2012. 81: p. 203–29.

3. Clague, M.J., S. Urbe, and D. Komander, Breaking the chains: deubiquitylating enzyme specificity begets function. Nat Rev Mol Cell Biol, 2019. 20(6): p. 338–352.

4. Hanna, J., D.S. Leggett, and D. Finley, Ubiquitin depletion as a key mediator of toxicity by translational inhibitors. Mol Cell Biol, 2003. 23(24): p. 9251–61.

5. Amerik, A., et al., In vivo disassembly of free polyubiquitin chains by yeast Ubp14 modulates rates of protein degradation by the proteasome. EMBO J, 1997. 16(16): p. 4826–38.

6. Kimura, Y. and K. Tanaka, Regulatory mechanisms involved in the control of ubiquitin homeostasis. J Biochem, 2010. 147(6): p. 793–8.

7. Kimura, Y., et al., An inhibitor of a deubiquitinating enzyme regulates ubiquitin homeostasis. Cell, 2009. 137(3): p. 549–59.

8. Stavreva, D.A., et al., Potential roles for ubiquitin and the proteasome during ribosome biogenesis. Mol Cell Biol, 2006. 26(13): p. 5131–45.

9. Vilotti, S., et al., The PML nuclear bodies-associated protein TTRAP regulates ribosome biogenesis in nucleolar cavities upon proteasome inhibition. Cell Death Differ, 2012. 19(3): p. 488–500.

10. Ounap, K., et al., The Stability of Ribosome Biogenesis Factor WBSCR22 Is Regulated by Interaction with TRMT112 via Ubiquitin-Proteasome Pathway. PLoS One, 2015. 10(7): p. e0133841.

11. Lopez, A.D., et al., Proteasomal degradation of Sfp1 contributes to the repression of ribosome biogenesis during starvation and is mediated by the proteasome activator Blm10. Mol Biol Cell, 2011. 22(5): p. 528–40.

12. Latonen, L., et al., Proteasome inhibitors induce nucleolar aggregation of proteasome target proteins and polyadenylated RNA by altering ubiquitin availability. Oncogene, 2011. 30(7): p. 790– 805.

13. Yasuda, S., et al., Stress- and ubiquitylation-dependent phase separation of the proteasome. Nature, 2020. 578(7794): p. 296–300.

14. Kong, K.E., et al., Timer-based proteomic profiling of the ubiquitin-proteasome system reveals a substrate receptor of the GID ubiquitin ligase. Mol Cell, 2021.

15. Silva, G.M., D. Finley, and C. Vogel, K63 polyubiquitination is a new modulator of the oxidative stress response. Nat Struct Mol Biol, 2015. 22(2): p. 116–23.

16. Kramer, G., A. Shiber, and B. Bukau, Mechanisms of Cotranslational Maturation of Newly Synthesized Proteins. Annu Rev Biochem, 2019. 88: p. 337–364.

17. Lafontaine, D.L. and D. Tollervey, The function and synthesis of ribosomes. Nat Rev Mol Cell Biol, 2001. 2(7): p. 514–20.

18. Woolford, J.L., Jr. and S.J. Baserga, Ribosome biogenesis in the yeast Saccharomyces cerevisiae. Genetics, 2013. 195(3): p. 643–81.

19. Mediani, L., et al., Defective ribosomal products challenge nuclear function by impairing nuclear condensate dynamics and immobilizing ubiquitin. EMBO J, 2019. 38(15): p. e101341.

20. Weids, A.J., et al., Distinct stress conditions result in aggregation of proteins with similar properties. Sci Rep, 2016. 6: p. 24554.

21. Cherkasov, V., et al., Systemic control of protein synthesis through sequestration of translation and ribosome biogenesis factors during severe heat stress. FEBS Lett, 2015. 589(23): p. 3654–64.

22. Koplin, A., et al., A dual function for chaperones SSB-RAC and the NAC nascent polypeptide-associated complex on ribosomes. J Cell Biol, 2010. 189(1): p. 57–68.

23. Sung, M.K., et al., A conserved quality-control pathway that mediates degradation of unassembled ribosomal proteins. Elife, 2016. 5.

24. Albert, B., et al., A ribosome assembly stress response regulates transcription to maintain proteome homeostasis. Elife, 2019. 8.

25. 25. 6hhhh Tye, B.W., et al., Proteotoxicity from aberrant ribosome biogenesis compromises cell fitness. Elife, 2019. 8.

26. Miller, S.B., et al., Compartment-specific aggregases direct distinct nuclear and cytoplasmic aggregate deposition. EMBO J, 2015. 34(6): p. 778–97.

27. Gallina, I., et al., Cmr1/WDR76 defines a nuclear genotoxic stress body linking genome integrity and protein quality control. Nat Commun, 2015. 6: p. 6533.

28. Mathew, V., et al., Selective aggregation of the splicing factor Hsh155 suppresses splicing upon genotoxic stress. J Cell Biol, 2017. 216(12): p. 4027–4040.

29. Martinez-Ferriz, A., et al., Ubiquitin-mediated mechanisms of translational control. Semin Cell Dev Biol, 2022. 132: p. 146–154.

30. Ikeuchi, K., et al., Collided ribosomes form a unique structural interface to induce Hel2-driven quality control pathways. EMBO J, 2019. 38(5).

31. Matsuki, Y., et al., Ribosomal protein S7 ubiquitination during ER stress in yeast is associated with selective mRNA translation and stress outcome. Sci Rep, 2020. 10(1): p. 19669.

32. Takehara, Y., et al., The ubiquitination-deubiquitination cycle on the ribosomal protein eS7A is crucial for efficient translation. iScience, 2021. 24(3): p. 102145.

33. Zhou, Y., et al., Structural impact of K63 ubiquitin on yeast translocating ribosomes under oxidative stress. Proc Natl Acad Sci U S A, 2020. 117(36): p. 22157–22166.

34. Ossareh-Nazari, B., et al., Ubiquitylation by the Ltn1 E3 ligase protects 60S ribosomes from starvation-induced selective autophagy. J Cell Biol, 2014. 204(6): p. 909–17.

35. Kraft, C., et al., Mature ribosomes are selectively degraded upon starvation by an autophagy pathway requiring the Ubp3p/Bre5p ubiquitin protease. Nat Cell Biol, 2008. 10(5): p. 602–10.

36. Blount, J.R., S.L. Johnson, and S.V. Todi, Unanchored Ubiquitin Chains, Revisited. Front Cell Dev Biol, 2020. 8: p. 582361.

37. Finley, D., et al., The ubiquitin-proteasome system of Saccharomyces cerevisiae. Genetics, 2012. 192(2): p. 319–60.

38. Piper, R.C., I. Dikic, and G.L. Lukacs, Ubiquitin-dependent sorting in endocytosis. Cold Spring Harb Perspect Biol, 2014. 6(1).

39. Xia, Z.P., et al., Direct activation of protein kinases by unanchored polyubiquitin chains. Nature, 2009. 461(7260): p. 114–9.

40. Hao, R., et al., Proteasomes activate aggresome disassembly and clearance by producing unanchored ubiquitin chains. Mol Cell, 2013. 51(6): p. 819–28.

41. Blount, J.R., et al., Isoleucine 44 Hydrophobic Patch Controls Toxicity of Unanchored, Linear Ubiquitin Chains through NF-kappaB Signaling. Cells, 2020. 9(6).

42. Clague, M.J., et al., Deubiquitylases from genes to organism. Physiol Rev, 2013. 93(3): p. 1289– 315.

43. Clague, M.J., C. Heride, and S. Urbe, The demographics of the ubiquitin system. Trends Cell Biol, 2015. 25(7): p. 417–26.

44. Costanzo, M., et al., A global genetic interaction network maps a wiring diagram of cellular function. Science, 2016. 353(6306).

45. Costanzo, M., et al., The genetic landscape of a cell. Science, 2010. 327(5964): p. 425–31.

46. Baryshnikova, A., et al., Quantitative analysis of fitness and genetic interactions in yeast on a genome scale. Nat Methods, 2010. 7(12): p. 1017–24.

47. Blount, J.R., et al., Expression and Regulation of Deubiquitinase-Resistant, Unanchored Ubiquitin Chains in Drosophila. Sci Rep, 2018. 8(1): p. 8513.

48. Blount, J.R., et al., Unanchored ubiquitin chains do not lead to marked alterations in gene expression in Drosophila melanogaster. Biol Open, 2019. 8(5).

49. Morimoto, D., et al., The unexpected role of polyubiquitin chains in the formation of fibrillar aggregates. Nat Commun, 2015. 6: p. 6116.

50. Yu, Y., et al., K29-linked ubiquitin signaling regulates proteotoxic stress response and cell cycle. Nat Chem Biol, 2021.

51. Meurer, M., et al., The regulatable MAL32 promoter in Saccharomyces cerevisiae: characteristics and tools to facilitate its use. Yeast, 2017. 34(1): p. 39–49.

52. Tong, A.H. and C. Boone, Synthetic genetic array analysis in Saccharomyces cerevisiae. Methods Mol Biol, 2006. 313: p. 171–92.

53. Wagih, O., et al., SGAtools: one-stop analysis and visualization of array-based genetic interaction screens. Nucleic Acids Res, 2013. 41(Web Server issue): p. W591-6.

54. den Brave, F., et al., Chaperone-Mediated Protein Disaggregation Triggers Proteolytic Clearance of Intra-nuclear Protein Inclusions. Cell Rep, 2020. 31(9): p. 107680.

55. Huh, W.K., et al., Global analysis of protein localization in budding yeast. Nature, 2003. 425(6959): p. 686–91.

56. Chong, Y.T., et al., Yeast Proteome Dynamics from Single Cell Imaging and Automated Analysis. Cell, 2015. 161(6): p. 1413–24.

57. Kraus, O.Z., et al., Automated analysis of high-content microscopy data with deep learning. Mol Syst Biol, 2017. 13(4): p. 924.

58. Ho, C.T., et al., Cellular sequestrases maintain basal Hsp70 capacity ensuring balanced proteostasis. Nat Commun, 2019. 10(1): p. 4851.

59. Kaganovich, D., R. Kopito, and J. Frydman, Misfolded proteins partition between two distinct quality control compartments. Nature, 2008. 454(7208): p. 1088–95.

60. Steinkraus, K.A., M. Kaeberlein, and B.K. Kennedy, Replicative aging in yeast: the means to the end. Annu Rev Cell Dev Biol, 2008. 24: p. 29–54.

61. Saarikangas, J. and Y. Barral, Protein aggregates are associated with replicative aging without compromising protein quality control. Elife, 2015. 4.

62. Martin-Villanueva, S., et al., Ubiquitin release from eL40 is required for cytoplasmic maturation and function of 60S ribosomal subunits in Saccharomyces cerevisiae. FEBS J, 2020. 287(2): p. 345– 360.

63. Lacombe, T., et al., Linear ubiquitin fusion to Rps31 and its subsequent cleavage are required for the efficient production and functional integrity of 40S ribosomal subunits. Mol Microbiol, 2009. 72(1): p. 69–84.

64. Finley, D., B. Bartel, and A. Varshavsky, The tails of ubiquitin precursors are ribosomal proteins whose fusion to ubiquitin facilitates ribosome biogenesis. Nature, 1989. 338(6214): p. 394-401.

65. McClellan, A.J., M.D. Scott, and J. Frydman, Folding and quality control of the VHL tumor suppressor proceed through distinct chaperone pathways. Cell, 2005. 121(5): p. 739–48.

66. Honma, Y., et al., TOR regulates late steps of ribosome maturation in the nucleoplasm via Nog1 in response to nutrients. EMBO J, 2006. 25(16): p. 3832–42.

67. Albanese, V., S. Reissmann, and J. Frydman, A ribosome-anchored chaperone network that facilitates eukaryotic ribosome biogenesis. J Cell Biol, 2010. 189(1): p. 69–81.

68. Meyer, A.E., L.A. Hoover, and E.A. Craig, The cytosolic J-protein, Jjj1, and Rei1 function in the removal of the pre-60 S subunit factor Arx1. J Biol Chem, 2010. 285(2): p. 961–8.

69. Loibl, M., et al., The drug diazaborine blocks ribosome biogenesis by inhibiting the AAA-ATPase Drg1. J Biol Chem, 2014. 289(7): p. 3913–22.

70. Prattes, M., et al., Structural basis for inhibition of the AAA-ATPase Drg1 by diazaborine. Nat Commun, 2021. 12(1): p. 3483.

71. Michel, M.A., et al., Assembly and specific recognition of k29- and k33-linked polyubiquitin. Mol Cell, 2015. 58(1): p. 95–109.

72. Morimoto, D. and M. Shirakawa, The evolving world of ubiquitin: transformed polyubiquitin chains. Biomol Concepts, 2016. 7(3): p. 157–67.

73. Braten, O., et al., Generation of free ubiquitin chains is up-regulated in stress and facilitated by the HECT domain ubiquitin ligases UFD4 and HUL5. Biochem J, 2012. 444(3): p. 611–7.

74. Hwang, C.S., et al., The N-end rule pathway is mediated by a complex of the RING-type Ubr1 and HECT-type Ufd4 ubiquitin ligases. Nat Cell Biol, 2010. 12(12): p. 1177–85.

75. Fang, N.N., et al., Hul5 HECT ubiquitin ligase plays a major role in the ubiquitylation and turnover of cytosolic misfolded proteins. Nat Cell Biol, 2011. 13(11): p. 1344–52.

76. Theodoraki, M.A., et al., A network of ubiquitin ligases is important for the dynamics of misfolded protein aggregates in yeast. J Biol Chem, 2012. 287(28): p. 23911–22.

77. Sitron, C.S. and O. Brandman, CAT tails drive degradation of stalled polypeptides on and off the ribosome. Nat Struct Mol Biol, 2019. 26(6): p. 450–459.

78. Panasenko, O.O. and M.A. Collart, Presence of Not5 and ubiquitinated Rps7A in polysome fractions depends upon the Not4 E3 ligase. Mol Microbiol, 2012. 83(3): p. 640–53.

79. Miller, S.B., A. Mogk, and B. Bukau, Spatially organized aggregation of misfolded proteins as cellular stress defense strategy. J Mol Biol, 2015. 427(7): p. 1564–74.

80. Tkach, J.M., et al., Dissecting DNA damage response pathways by analysing protein localization and abundance changes during DNA replication stress. Nat Cell Biol, 2012. 14(9): p. 966–76.

81. Mathew, V., et al., Cdc48 regulates intranuclear quality control sequestration of the Hsh155 splicing factor in budding yeast. J Cell Sci, 2020. 133(23).

82. Ghosh, A., et al., Proteotoxic stress promotes entrapment of ribosomes and misfolded proteins in a shared cytosolic compartment. Nucleic Acids Res, 2020. 48(7): p. 3888–3905.

83. Pathak, B.K., et al., Sequestration of Ribosome during Protein Aggregate Formation: Contribution of ribosomal RNA. Sci Rep, 2017. 7: p. 42017.

84. Nanduri, P., et al., Chaperone-mediated 26S proteasome remodeling facilitates free K63 ubiquitin chain production and aggresome clearance. J Biol Chem, 2015. 290(15): p. 9455–64.

85. Ouyang, H., et al., Protein aggregates are recruited to aggresome by histone deacetylase 6 via unanchored ubiquitin C termini. J Biol Chem, 2012. 287(4): p. 2317–27.

86. Du, M., et al., Liquid phase separation of NEMO induced by polyubiquitin chains activates NF-kappaB. Mol Cell, 2022.

87. Baker, N.E., Emerging mechanisms of cell competition. Nat Rev Genet, 2020. 21(11): p. 683–697.

88. Baumgartner, M.E., et al., Proteotoxic stress is a driver of the loser status and cell competition. Nat Cell Biol, 2021. 23(2): p. 136–146.

89. Recasens-Alvarez, C., et al., Ribosomopathy-associated mutations cause proteotoxic stress that is alleviated by TOR inhibition. Nat Cell Biol, 2021. 23(2): p. 127–135.

90. Ogrodnik, M., et al., Dynamic JUNQ inclusion bodies are asymmetrically inherited in mammalian cell lines through the asymmetric partitioning of vimentin. Proc Natl Acad Sci U S A, 2014. 111(22): p. 8049–54.

91. Aguilaniu, H., et al., Asymmetric inheritance of oxidatively damaged proteins during cytokinesis. Science, 2003. 299(5613): p. 1751–3.

92. Winston, F., C. Dollard, and S.L. Ricupero-Hovasse, Construction of a set of convenient Saccharomyces cerevisiae strains that are isogenic to S288C. Yeast, 1995. 11(1): p. 53–5.

93. Brachmann, C.B., et al., Designer deletion strains derived from Saccharomyces cerevisiae S288C: a useful set of strains and plasmids for PCR-mediated gene disruption and other applications. Yeast, 1998. 14(2): p. 115–32.

94. Janke, C., et al., A versatile toolbox for PCR-based tagging of yeast genes: new fluorescent proteins, more markers and promoter substitution cassettes. Yeast, 2004. 21(11): p. 947–62.

95. Gietz, R.D. and R.A. Woods, Transformation of yeast by lithium acetate/single-stranded carrier DNA/polyethylene glycol method. Methods Enzymol, 2002. 350: p. 87–96.

96. Chawade, A., E. Alexandersson, and F. Levander, Normalyzer: a tool for rapid evaluation of normalization methods for omics data sets. J Proteome Res, 2014. 13(6): p. 3114–20.

97. Tollervey, D., A yeast small nuclear RNA is required for normal processing of pre-ribosomal RNA. EMBO J, 1987. 6(13): p. 4169–75.

98. Sambrook, J. and D.W. Russell, Separation of RNA According to Size: Electrophoresis of Glyoxylated RNA through Agarose Gels. CSH Protoc, 2006. 2006(1).

99. David, F.P., et al., HTSstation: a web application and open-access libraries for high-throughput sequencing data analysis. PLoS One, 2014. 9(1): p. e85879.

